# STUB1 is an intracellular checkpoint for interferon gamma sensing

**DOI:** 10.1101/2020.12.14.420539

**Authors:** Simon Ng, Shuhui Lim, Adrian Chong Nyi Sim, Ruban Mangadu, Ally Lau, Chunsheng Zhang, Sarah Bollinger Martinez, Arun Chandramohan, U-Ming Lim, Samantha Shu Wen Ho, Shih Chieh Chang, Pooja Gopal, Lewis Z. Hong, Adam Schwaid, Aaron Zefrin Fernandis, Andrey Loboda, Cai Li, Uyen Phan, Brian Henry, Anthony W. Partridge

**Author notes:** Corresponding author: Anthony W. Partridge.

## Abstract

Immune checkpoint blockade (ICB) leads to durable and complete tumour regression in some patients but in others gives temporary, partial or no response. Accordingly, significant efforts are underway to identify tumour-intrinsic mechanisms underlying ICB resistance.

Results from a published CRISPR screen in a mouse model suggested that targeting STUB1, an E3 ligase involved in protein homeostasis, may overcome ICB resistance but the molecular basis of this effect remains unclear. Herein, we report an under-appreciated role of STUB1 to dampen the interferon gamma (IFNγ) response. Genetic deletion of STUB1 increased IFNGR1 abundance on the cell surface and thus enhanced the downstream IFNγ response as showed by multiple approaches including Western blotting, flow cytometry, qPCR, phospho-STAT1 assay, immunopeptidomics, proteomics, and gene expression profiling. Human prostate and breast cancer cells with STUB1 deletion were also susceptible to cytokine-induced growth inhibition.

Furthermore, blockade of STUB1 protein function recapitulated the *STUB1*-null phenotypes. Despite these encouraging *in vitro* data and positive implications from clinical datasets, we did not observe *in vivo* benefits of inactivating *Stub1* in mouse syngeneic tumour models – with or without combination with anti-PD-1 therapy. However, our findings elucidate STUB1 as a barrier to IFNγ sensing, prompting further investigations to assess if broader inactivation of human STUB1 in both tumors and immune cells could overcome ICB resistance.

## Introduction

Immune checkpoint blockade (ICB) unleashes the adaptive immune system to fight cancer and results in long-term patient survival unmatched by other drug treatments. Substantial evidence has highlighted IFNγ response and antigen presentation as key components for cancer immunosurveillance and immunotherapy^1–11^. As an emerging paradigm^1,12–14^, intact IFNγ sensing in tumours leads to adequate antigen presentation and T cell recognition, but also upregulates programmed death ligand 1 (PD-L1) to confer immunoevasion^15^. Blockade of PD-1/PD-L1 in these patients reinvigorated anti-tumour activity of exhausted T cells and resulted in durable tumour regression^16–18^. In contrast, patients with poor anti-PD-1 response have low tumour-infiltrating lymphocytes, low expression of PD-L1 and reduced antigen presentation^19–21^. These ICB-resistant tumours have either preexisting, or post-treatment acquired resistance caused by defective interferon signaling^4,5,22^, reduced sensitivity to IFNγ^4,20,23–25^, or attenuated antigen presentation not explained by disruptive pathway mutations^21,26–30^. In these circumstances, treating IFNγ-insensitive tumours with ICB is ineffective, thus demanding different approaches or a combination with ICB.

Several research groups have employed genetic loss-of-function screens to identify targets that underly ICB resistance or the targets needed for anti-tumour immunity. By mining the repository of CRISPR screens (BioGRID ORCS)^31^, we realized that loss of *Stub1* appears to reverse the resistance of immunotherapy in an *in vivo* tumour mouse model^9^ and enhance *in vitro* T cell-mediated killing of murine tumour cells, e.g., B16-F10^10^, CT26^32^ and Renca^32^. Furthermore, low STUB1 correlates with high PD-L1 in human HAP1 and A375 cells^33^. Although the underlying mechanisms remain elusive, recurring discovery of STUB1 among the top 1% hits in these genetic screens, as summarized in Supplementary Table 1, highlight a significant, yet under-appreciated role of STUB1 in regulating anti-tumour immunity. Based on the unbiased screening results and the canonical role of STUB1 in response to stress stimuli^34–36^, we hypothesize that STUB1 may play a conserved and prominent part in dampening stress triggered by the immune system.

To study the molecular role of *Stub1*, we used virus-free CRISPR-editing to delete *Stub1* from an ICB-resistant and poorly immunogenic^37^ murine melanoma line (B16-F10) by electroporating the corresponding crRNA/tracrRNA/Cas9 ribonucleoprotein (RNP) into the cells. This transient approach is favoured as recent reports have highlighted the capability of viral based approaches to artificially increase immunogenicity of syngeneic mouse lines^38,39^. After CRISPR editing, we isolated a total of nine *Stub1*-null cells by single-cell subcloning (Supplementary Table 2). For most experiments, we focused on two clonal cell lines – gStub1 #1 (1D10) and gStub1 #2 (1A12) – targeted by two independent crRNAs respectively.

### *Stub1* deficiency enhances antigen presentation *via* increased IFN**γ** responsiveness

Tumours often reduce antigen presentation to evade immunosurveillance and immunotherapy^5,21,27,40^. Accordingly, we measured the effect of *Stub1* deletion on major histocompatibility complex class I (MHC-I) surface expression in B16-F10 cells by flow cytometry. Strikingly, compared to parental and control cells, tumour cells lacking *Stub1* displayed significantly higher, IFNγ-dependent, MHC-I on cell surface (Fig. 1a). The differential antigen presentation is consistently found across all nine *Stub1*-null cells isolated *via* single-cell subcloning (Supplementary Fig. 1a-e and Supplementary Table 2). We regularly saw differential expression of MHC-I at different doses of IFNγ (Fig. 1b–c and Supplementary Fig. 1f). The IFNγ-STAT1-IRF1 axis induces genes associated with antigen processing and presentation.

**Fig. 1.**
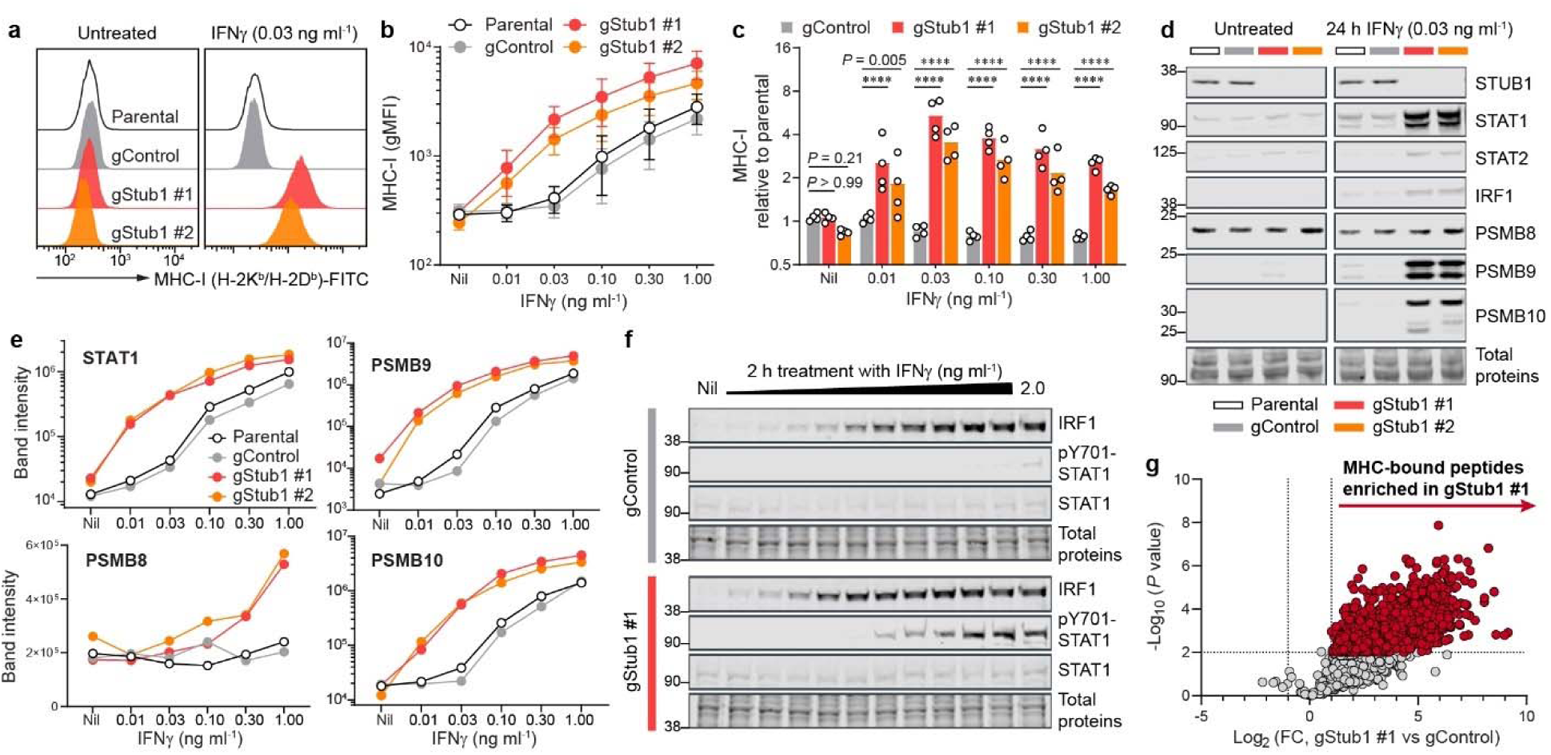
*Stub1* deletion enhances antigen processing and presentation by sensitizing tumour cells to IFNγ. **a**–**c**, Flow cytometry analysis of cell surface MHC-I on parental, control or independent *Stub1*-null B16-F10 cells. gMFI, geometric mean fluorescence intensity. **d**, **e**, Western blot analysis of the expression level of STUB1, STAT1, STAT2, IRF1, PSMB8, PSMB9 and PSMB10 in parental, control or independent *Stub1*-null B16-F10 cells. Band intensity was normalized with total protein signal. The tumour cells were either untreated (Nil) or treated with IFNγ for 24 h (a–e). See Supplementary Fig. 1 for additional flow cytometry plots, Western blot data and analysis (a, d, e). **f**, Western blot analysis of the expression level of IRF1, STAT1, and phosphorylation of Tyr701-STAT1 at 2 h post-treatment with IFNγ (2-fold serial dilution from 2.0 ng ml^−1^). **g**, Volcano plot showing differential presentation of MHC-associated peptide in gStub1 #1 versus gControl cells, following stimulation with 0.03 ng ml^−1^ IFNγ for 24 h. Red circles highlight peptides significantly enriched in gStub1 #1 cells (2-fold cutoff, *P* ≤0.01; *n* = 3 biological replicates). FC, fold change. See Supplementary Fig. 2d for data of gStub1 #2 cells. Representative of four (a) or two (d–f) independent experiments. Data are mean ± s.d. (b) or mean with all data points (c) from four independent experiments. *P* values were determined by ordinary two-way ANOVA on Log2-transformed data with Dunnett’s multiple comparisons test, **** *P* ≤0.0001 (c).

Remarkably, *Stub1* deletion led to upregulation of STAT1, STAT2, and IRF1 after 24 h stimulation with IFNγ (Fig. 1d). Immunoproteasome complex has been associated with better tumour immunogenicity and better prognosis and response to checkpoint therapies in melanoma^41^. Similarly, in an IFNγ-dependent manner, *Stub1* deletion upregulated PSMB8, PSMB9 and PSMB10 – subunits of the immunoproteasome complex (Fig. 1d–e). The differential protein expression of STAT1, STAT2, IRF1, PSMB9 and PSMB10 are maintained across all doses of IFNγ, while PSMB8 upregulation is more pronounced at doses higher than 0.30 ng ml^−1^ (Fig. 1e and Supplementary Fig. 1g–h). To measure the acute response of the signal transduction, we treated the tumour cells with a titration of IFNγ and harvested the cellular lysates for analysis at 2 h post-stimulation. Loss of *Stub1* lowers the stimulating threshold of IFNγ needed for the early induction of IRF1 and the phosphorylation of Tyr701-STAT1 (Fig. 1f). Total STAT1 protein level stayed low and stable during the initial response (Fig. 1f) but was significantly upregulated at 24 h post-stimulation (Fig. 1d). To probe the diversity of the MHC-associated peptides presented on the tumour cells, we immunoprecipitated the MHC-I to identify and quantify MHC-bound peptides with mass spectrometry (Supplementary Fig. 2a–c).

Immunopeptidomic analysis definitively confirmed a global upregulation of antigen presentation on *Stub1*-null cells relative to the control line (Fig. 1g and Supplementary Fig. 2d). Overall, parental B16-F10 and control cells demanded at least 10-fold higher concentration of IFNγ to achieve a comparable response seen in *Stub1*-null cells (Fig. 1b and 1e), suggesting *Stub1* is a key checkpoint for IFNγ sensing in tumour cells.

### Stub1 constrains IFNγ response by downregulating IFNγ receptors

To investigate how *Stub1* constitutively suppresses the IFNγ response, we probed the level of IFNγ receptor 1 (IFNGR1) in B16-F10 cells (Fig. 2a). Indeed, loss of *Stub1* increased the surface expression level of IFNGR1 under both resting and IFNγ-stimulating conditions (Fig. 2b–c and Supplementary Fig. 3a). Interestingly, the cell surface level of IFNGR1 declined with increasing IFNγ concentration (Fig. 2b), perhaps through feedback endocytosis of the ligand-receptor complexes^42^. The regulation was specific as *Stub1* deletion had no significant effect on other cytokine receptors, such as IL1R1, IL6R, GP130 and IFNAR1 (Fig. 2d, Supplementary Fig. 3b–c). Stable gene expression of *Ifngr1* suggested that downregulation of the receptor by the E3 ligase STUB1 occurs at the protein level (Supplementary Fig. 3d).

**Fig. 2.**
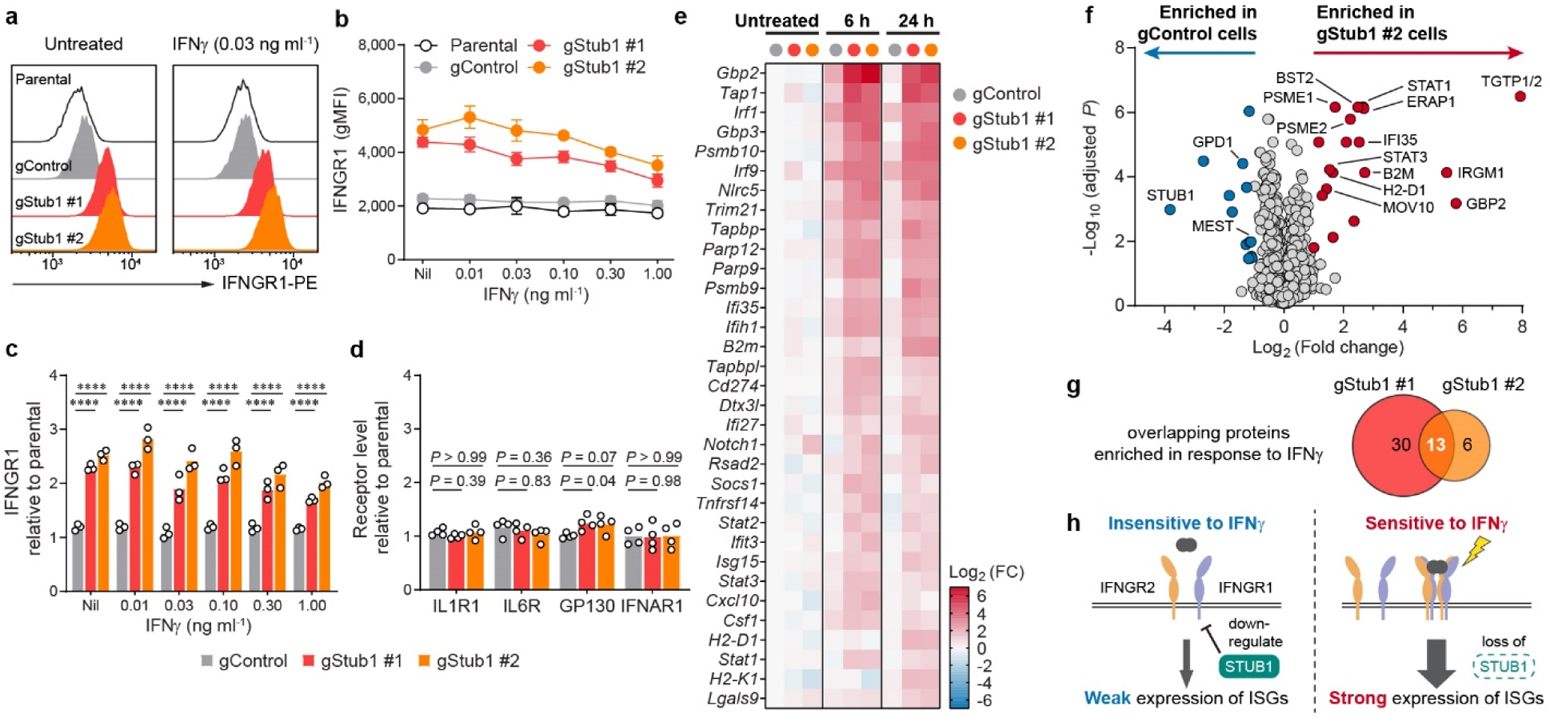
*Stub1* dampens IFNγ sensing by downregulating IFNGR1. **a**–**c**, Flow cytometry analysis of cell surface IFNGR1 on parental, control or independent *Stub1*-null B16-F10 cells which were either untreated (Nil) or treated with IFNγ for 24 h. See Supplementary Fig. 3a for additional plots. **d**, Flow cytometry analysis of the surface level of other cytokine receptors on the tumour cells. See Supplementary Fig. 3b–c for the plots. **e**, Heatmap showing genes (Supplementary Table 3) being upregulated by >2-fold in both gStub1 #1 and #2 cells relative to untreated gControl cells. The cells were treated with 0.03 ng ml^−1^ IFNγ for 6 or 24 h. See Supplementary Fig. 4a for the full heatmap. FC, fold change. **f**, Volcano plot showing differential protein expression in gStub1 #2 versus gControl cells, following stimulation with 0.03 ng ml^−1^ IFNγ for 24 h. Red or blue circles highlight proteins significantly enriched in gStub1 #2 or gControl cells respectively (2-fold cutoff, adjusted *P* ≤0.05; *n* = 6 replicates per cell group, 3 biological replicates × 2 MS replicates). See Supplementary Fig. 4e for data of gStub1 #1 cell. **g**, MS proteomics uncovered 13 proteins commonly enriched in both gStub1 #1 and #2 cells. The overlapping proteins are explicitly labeled in panel **f**. **h**, Proposed model whereby *Stub1* is an intracellular checkpoint that curbs the tumour cells’ ability to sense and respond to IFNγ by downregulating IFNGR1. Representative of three independent experiments (a). Data are mean ± s.d. (b) or mean with all data points (c) from three independent experiments. Data are mean with all data points from four independent experiments (d). *P* values were determined by ordinary two-way ANOVA (c) or one-way ANOVA (d) on Log2-transformed data with Dunnett’s multiple comparisons test, **** *P* ≤0.0001.

We reasoned that constitutive upregulation of IFNGR1 in *Stub1*-null cells could potentiate and amplify downstream signal transduction. To broadly evaluate the response, we studied the gene expression of 750 immune-related genes (NanoString PanCancer IO 360; Supplementary Table 3). Overall, most genes have comparable expression among the *Stub1*-null and control cells (Supplementary Fig. 4a). As expected, *Stub1*-null cells had an enhanced response to IFNγ treatment (6 h or 24 h) as evidence by the increased induction of interferon-stimulated genes (ISGs), including those that govern the interferon signaling pathway (*Stat1*, *Stat2*, *Irf1* and *Irf9*), antigen processing and presentation (*H2-D1*, *H2-K1*, *B2m*, *Nlrc5*, *Tap1*, *Tapbp*, *Tapbpl*, *Psmb9* and *Psmb10*), and chemotaxis of immune cells (*Cxcl10* and *Csf1*) (Fig. 2e, Supplementary Fig. 4b–c). In contrast, the control cells weakly induced these ISGs, in response to stimulation with a low dose of IFNγ for 6 h, and the ISGs mostly receded at 24 h post-stimulation (Fig. 2e, Supplementary Fig. 4d). Importantly, *Stub1* did not directly regulate the ISGs themselves, as evidenced by their comparable gene expression (Fig. 2e and Supplementary Fig. 3d) and protein levels (Fig. 1d–f) in the untreated *Stub1*-null and control cells.

To investigate IFNγ signaling at the protein level, we performed proteome-wide analysis with mass spectrometry (MS). A high-quality dataset consisting of ∼2300 proteins (Supplementary Table 4) definitively validated our hypothesis – STUB1 is a checkpoint and barrier for IFNγ sensing. Loss of STUB1 sensitized tumour cells to IFNγ exposure and led to statistically significant enrichment of the protein targets of ISGs, including those required for antigen presentation such as H2-K1, B2M, PSME1, PSME2 and ERAP1 (Fig. 2f). Overall, we identified an overlapping set of 13 proteins (explicitly labeled in Fig. 2f), all inducible by interferon, being enriched in both independent *Stub1*-null cells relative to the control cells (Fig. 2g and Supplementary Fig. 4e). Taken together, we propose a framework whereby STUB1 may confer ICB resistance by downregulating IFNGR1 on the cell surface, thus curbing the tumour cells’ ability to sense and respond to IFNγ (Fig. 2h).

### Inhibition of STUB1 phenocopies the genetic knockout

A recent study^43^ identified a high-affinity peptide (SIWWPD) capable of blocking the interaction of STUB1 with HSPA8 – a chaperone bound to STUB1 through its C-terminal peptide. We validated the binding of the inhibitory peptide using multiple orthogonal biophysical assays^44^, including isothermal titration calorimetry (K_D_ = 14 ± 2 nM, Fig. 3a and Supplementary Fig. 5a–c), thermal shift assay (⊗T_m_ = 18.3 ± 0.1 °C, Fig. 3b), and competitive fluorescence polarization assay (IC_50_ = 0.34 ± 0.02 μM, Fig. 3c). We also designed a control peptide (SIWWHR), where STUB1 binding is abolished (K_D_ > 10 μM, IC_50_ > 100 μM, ⊗T_m_ = −0.1 ± 0.2 °C, Fig. 3a–c and Supplementary Fig. 5c–d) by substituting two key interacting residues (Pro-Asp) with counter-productive ones (His-Arg). To investigate if stoichiometric STUB1 inhibition could recapitulate the *Stub1*-null phenotypes, we engineered B16-F10 cells to constitutively and stably express a fusion protein consisting of an mCherry2 reporter^45^ tagged on its C-terminus with the inhibitory peptide or control sequence (Fig. 3d–e). As expected, ectopic expression of mCherry2-SIWWPD, but not its control, led to upregulation of IFNGR1 on the cell surface of B16-F10, under both resting and IFNγ-stimulating conditions (Fig. 3f). Furthermore, this effect was not restricted to murine cells as stable expression of the inhibitory biologic in human tumour cells (A375 and A549) also resulted in the same phenotype (Fig. 3f), which in turn potentiated the cells to boost the surface levels of MHC-I in response to IFNγ (Fig. 3g). Importantly, the interaction between the expressed biologic and STUB1 is specific, as STUB1 was co-precipitated with FLAG-mCherry2-SIWWPD, but not its control, from the cellular lysate (Fig. 3h). The interaction is completely reversible in a dose-dependent manner by spiking synthetic peptide inhibitor into the mixture of co-immunoprecipitation. Overall, stoichiometric inhibition of STUB1 with the expressed biologic completely recapitulated the phenotypes of *Stub1*-null cells shown earlier (Fig. 1c for MHC-I, and Fig. 2c for IFNGR1), an important result as pharmacological inhibition may not always mimic the outcome of a genetic knockout.

**Fig. 3.**
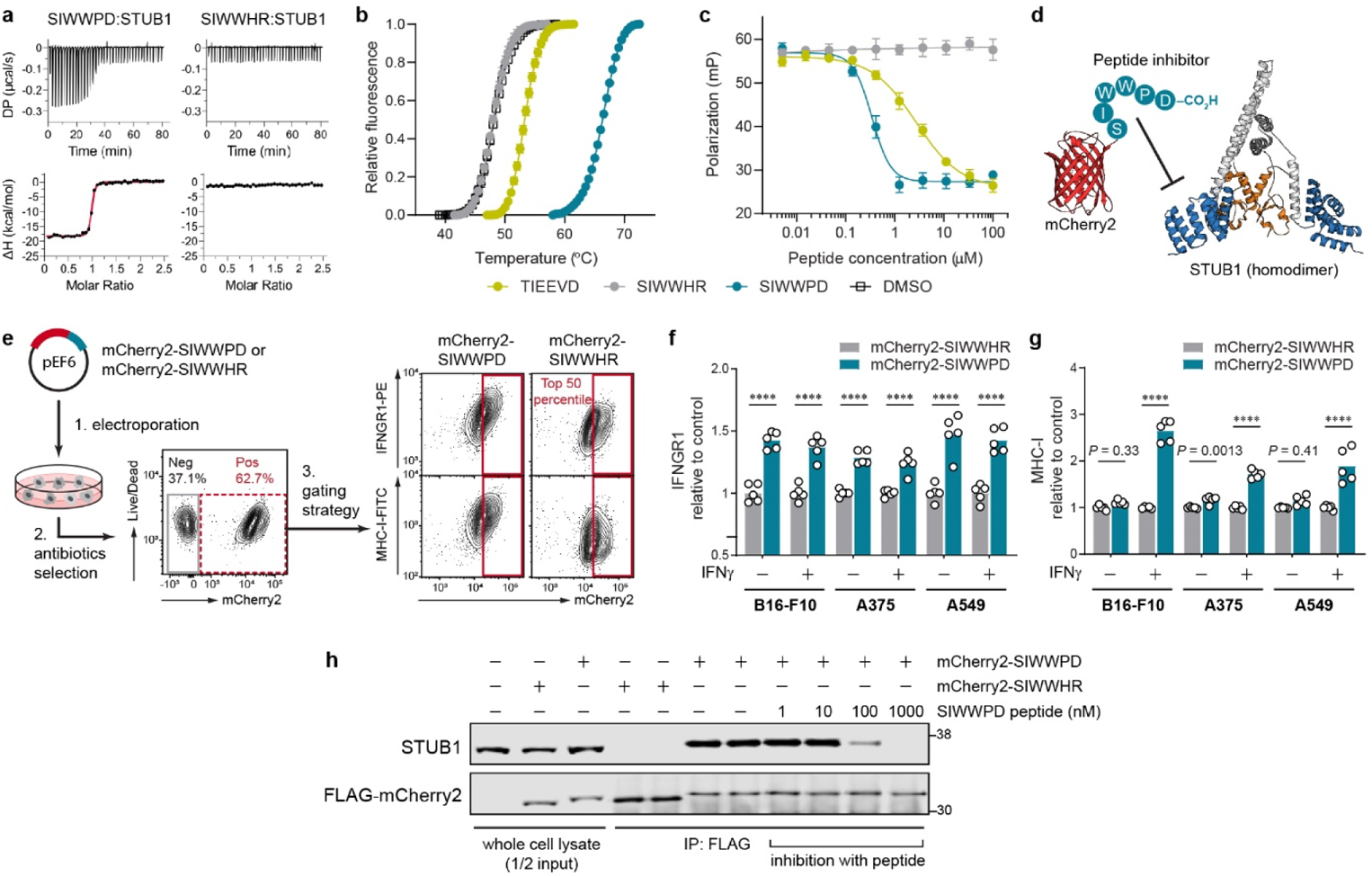
Pharmacological inhibition of STUB1 with expressed biologic phenocopies the genetic knockout. **a**, **b**, Validation of the binding of synthetic peptides to STUB1 (aa25–aa153) by isothermal titration calorimetry (a) and thermal shift assay (b). Representative of three independent experiments (a). Data are mean ± s.d. of six replicates derived from three independent experiments (b). **c**, Competitive fluorescence polarization assay. Synthetic peptides were assessed for their ability to compete with 15 nM of tracer peptide (5-FAM-SSGPTIEEVD) for binding to 1 μM STUB1 (aa25–aa153). Data are mean ± s.d. of six replicates derived from two independent experiments. **d**, Design of the inhibitory biologic by grafting the peptide (SIWWPD) to the C-terminus of an mCherry2 (red) scaffold. The fused peptide blocks the function of the tetratricopeptide repeat domain (blue) of STUB1 (PDB code 2C2L) and inhibits its substrate binding. U-box domain (orange) which recruits the E2 ubiquitin-conjugating enzyme is not affected. **e**, Generation of tumour cell lines stably expressing the biologic or its control. Plasmid encoding the biologic was electroporated into tumour cells, followed by antibiotics selection of the stable clones. The mCherry2-positive cells (red dotted box) were further gated for mCherry2^hi^ population (top 50th percentile, red box). Gating example represents IFNγ-treated B16-F10 stable cell lines. **f**, **g**, Flow cytometry analysis of the relative cell surface level of IFNGR1 (f) and MHC-I (g) expressed by the mCherry2^hi^ population in B16-F10, A375 or A549 cells. The cells were either untreated or treated with mouse IFNγ (0.03 ng ml^−1^) or human IFNγ (0.01 ng ml^−1^) for 24 h. The expression levels were normalized to the average value of the control (mCherry2-SIWWHR). *n* = 5 biological replicates from two independent experiments (f–g). Bars are mean with all data points (f–g). *P* values were determined by ordinary two-way ANOVA in each cell type with Sidak’s multiple comparisons test, **** *P* ≤0.0001 (f–g). **h**, Co-immunoprecipitation (co-IP) of FLAG-mCherry2-peptide and STUB1 from the cellular lysate of B16-F10 using anti-FLAG antibody. Synthetic peptide (SIWWPD) was added into the co-IP mixture to assess specificity of the interaction. Blot is representative of three independent experiments.

### Clinical relevance of *STUB1* across multiple tumours

Earlier analysis of data from KEYNOTE clinical trials showed that tumour mutational burden (TMB) and an 18-gene T-cell inflamed, IFNγ-related gene expression profile (GEP) has predictive value in identifying anti-PD-1 responders and non-responders^7,18^. TMB and GEP have low correlation and are tissue-agnostic measures that independently predict anti-PD-1 responsiveness in multiple tumours. So, we analyzed the correlation of *STUB1* with TMB and GEP using the bulk RNAseq data from The Cancer Genome Atlas (TCGA) dataset. The resulting analysis showed that *STUB1* is slightly depleted in tumours associated with high GEP score (top 55th percentile, GEP^hi^) regardless of the TMB value (Fig. 4a), suggesting low *STUB1* correlates with inflamed tumour microenvironment. To explore the expression level of *STUB1* in different cell types, we deconvoluted the bulk RNAseq data with CIBERSORT analysis^46^ in each tumour type from TCGA (Fig. 4b). In general, *STUB1* is low in the immune effector cells, such as activated NK cells, CD8^+^ T cell, γδ^+^ T cell, and activated dendritic cells. Interestingly, *STUB1* is enriched in M0 and M2 macrophage, compared to M1 macrophage, across most of the tumour types. These analyses were repeated using the Moffit dataset (Supplementary Fig. 6a, 6b) and the trends are mostly consistent with the TCGA dataset (Fig. 4a–b). Finally, we compared *STUB1* expression in tumours and adjacent normal tissues across multiple tumour types for which the data are available in TCGA (Fig. 4c). *STUB1* is overexpressed in thyroid, kidney, prostate, and breast tumours compared to their adjacent normal tissues, while a reverse trend is found in gastric cancer. Overall, the association of underexpression of *STUB1* in an inflamed tumour microenvironment (GEP^hi^) and the overexpression of *STUB1* in immunologically “cold” tumours (prostate and breast) support our interpretation of *STUB1* as an immunosuppressive gene, which likely constrains IFNγ sensing in the cancer-immunity cycle^47^.

**Fig. 4.**
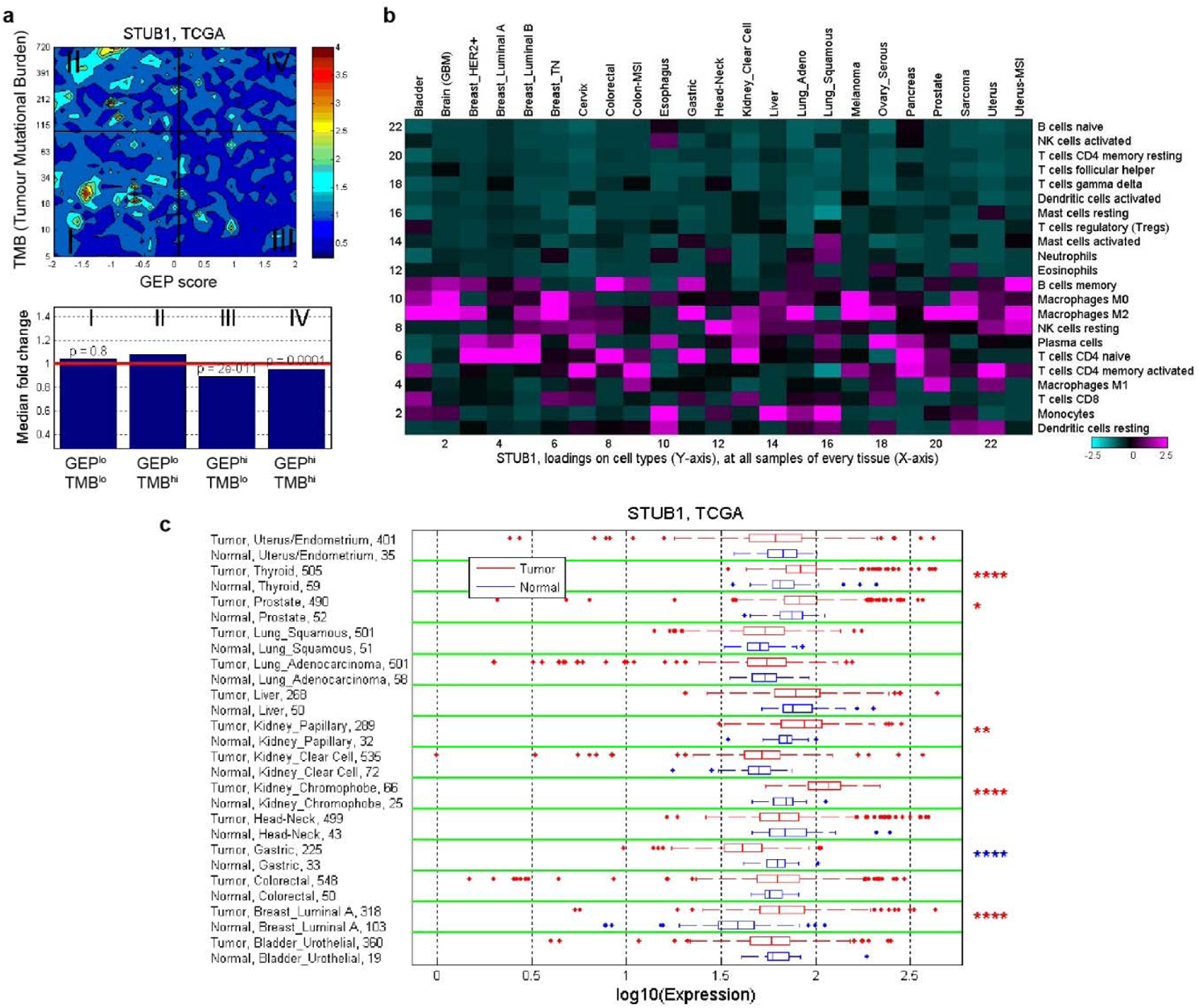
Correlation and expression of *STUB1* gene in TCGA dataset. **a**, Contour plot illustrates the association of *STUB1* with TMB and GEP. Blue and red represent under- and overexpression, respectively. TMB cut-off was set at 100 and GEP cut-off corresponds to 55th percentile value for pan-cancer cohort. **b**, *In-silico* deconvolution analysis of bulk RNAseq data from TCGA was used to establish the association between *STUB1* expression and different cell types. Deconvolution analysis was performed separately for each tumor type. **c**, Expression of *STUB1* in tumor tissue and adjacent normal tissue is compared across tumor types for which both tumor and adjacent normal samples are available in TCGA dataset. The significance of the difference is indicated with * *P* ≤0.05, ** *P* ≤0.01, and **** *P* ≤0.0001.

### *STUB1* deletion sensitized tumour cells to growth inhibition induced by cytokines

Next, we asked if inactivation of *STUB1* could sensitize human tumour cells to growth inhibition induced by the cytokines such as IFNγ and TNFα. We shortlisted DU145 (prostate), PC-3 (prostate) and MCF7 (ER+/HER2-breast) as model cell lines for the study, due to the overexpression of *STUB1* in prostate and breast cancer suggested by analysis of the TCGA dataset (Fig. 4c). As a benchmark, we also inactivated *PTPN2*, a well-studied negative regulator of IFN signaling, in parallel. Delivery of sgRNA/Cas9 RNP complexes using electroporation resulted in near complete knockout (>99%) of *STUB1* in all three tumour lines. In addition, we achieved sufficient knockout of *PTPN2* in DU145 (96%), PC-3 (83%) and MCF7 (89%) as determined by Western blot (Fig. 5a). Importantly, loss of *STUB1*, but not *PTPN2*, increased the surface expression level of IFNGR1 in all three tumour lines (Fig. 5b), a result highly consistent with B16-F10 cells (Fig. 2a). By measuring the phosphorylation of Tyr701-STAT1 as a proxy of signaling activation, we confirmed that loss of either *PTPN2* or *STUB1* increased the cells’ response to IFNγ in a dose-dependent manner (Fig. 5c). Generally, *PTPN2* is a stronger negative regulator as compared to *STUB1*, except for MCF7 in which deletion of *PTPN2* or *STUB1* enhanced the signaling to a similar extent. Prolonged treatment with IFNγ could inhibit the growth of tumour cells.^48^ Indeed, treatment with IFNγ alone or combined IFNγ and TNFα, but not TNFα alone, sensitized *PTPN2*- or *STUB1*-null prostate tumour cells to growth inhibition *via* measuring the ATP level produced by viable cells (Fig. 5d). Surprisingly, inactivation of *PTPN2* or *STUB1* could sensitize MCF7 breast tumour cell, a TNFα-sensitive line^49^, to growth inhibition induced by TNFα alone, albeit to a smaller extent as compared with inhibition induced by IFNγ or its combination with TNFα. To exclude the possibility that the genetic knockout could affect the ATP level in unexpected ways, we measured the live-cell protease activity as a surrogate for cell viability. This orthogonal assay confirmed that the antiproliferative effects and trends (Supplementary Fig. 7) are reproducible and highly consistent with the results obtained by ATP assay (Fig. 5d).

**Fig. 5.**
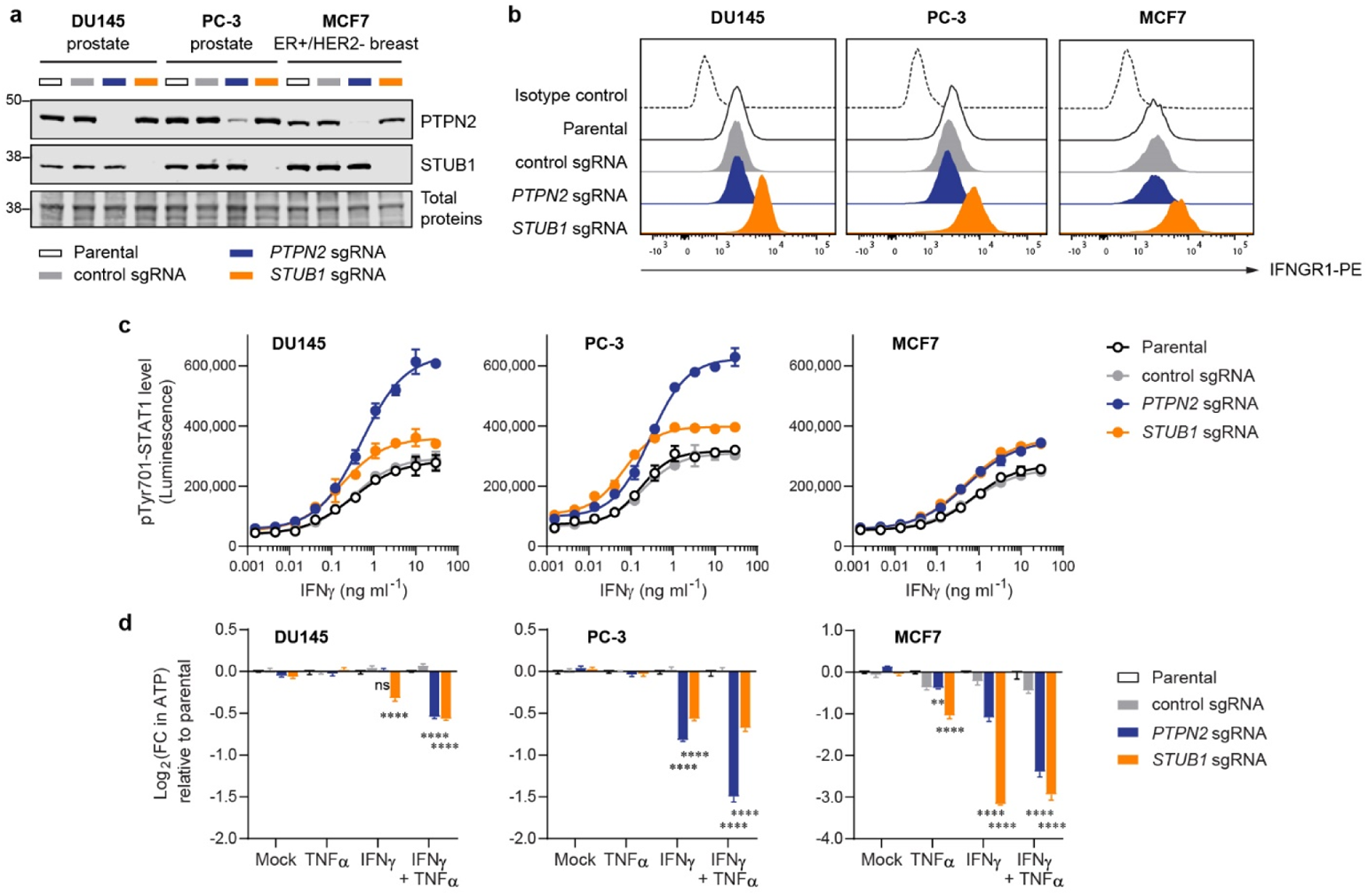
Inactivation of *STUB1* or *PTPN2* sensitized human tumour cells to growth inhibition induced by cytokines. **a**, Western blot analysis of the expression level of PTPN2 and STUB1 in tumour cells. **b**, Flow cytometry plot showing the surface expression level of IFNGR1 in tumour cells. In parallel, parental cells were stained with PE-conjugated isotype control antibody to demonstrate low level of non-specific binding. **c**, Level of phosphorylated Tyr701-STAT1 after 30-min response to varying doses of IFNγ as measured by Lumit immunoassay. **d**, Fold change (FC) in ATP level relative to the parental cells as a quantification of viable cells. Measurements were performed using CellTiter-Glo 2.0 assay after 6-day treatment with the cytokines (10 ng ml^−1^ each). Data are mean ± s.d. from two biological replicates (c) or mean ± s.e.m. from three biological replicates (d). *P* values were determined by ordinary two-way ANOVA on Log2-transformed data with Dunnett’s multiple comparisons test versus parental cells, ** *P* ≤0.01, **** *P* ≤0.0001, ns *P* >0.90 (d). Representative of two independent experiments (a–d).

### *Stub1* deletion provided limited benefits in syngeneic mouse models

To understand if inactivation of *Stub1* could sensitize tumours to immunotherapy, we inoculated the CRISPR-edited B16-F10 clonal cells into immuno-competent C57BL/6J mice. We treated the mice with GVAX vaccine (GM-CSF secreting, irradiated B16-F10 cells), followed by anti-PD-1 antibody (Fig. 6a). However, we observed no benefit by inactivating *Stub1* in the transplanted tumour cells (Fig. 6b). Compared to the control, gStub1 #1 clonal cells formed more aggressive tumours, though not statistically significant (Fig. 6b, *P* = 0.10 at day 16). Survival analysis suggested that tumours from gStub1 #1 were indeed more aggressive (Fig. 6c). Mice bearing the gStub1 #1 tumours demonstrated shorter median survival (22 days, *P* = 0.007) as compared to the control group (28 days). In contrast, 4 out of 15 mice grafted with gStub1 #2 clonal cells achieved complete tumour regression, while only 1 out of 15 mice grafted with the control cells had a complete response and none of the mice grafted with gStub1 #1 cells survived. The conflicting results between gStub1 #1 and gStub1 #2 are expected to be contributed by the intrinsic differences between the clonal cells (Fig. 2g, Supplementary Fig. 2e–f). To circumvent the confounding effect, we selected a different model, CT26 colon tumour cells, for further studies. We transiently delivered sgRNA/Cas9 RNP complexes to the tumour cells using electroporation (a virus-free approach). Instead of single-cell subcloning, we sorted the population for high expression of IFNGR1 to enrich the *Stub1*-null cells (Supplementary Fig. 9a). The phenotypes, namely the relative expression level of IFNGR1, MHC-I, STAT1, IRF1 and PSMB9 of the *Stub1*-null vs control CT26 cells (Supplementary Fig. 9b–i), were highly consistent with previous results obtained for B16-F10 (Fig. 1 and Fig. 2). We then transplanted the CRISPR-edited CT26 cells into immuno-competent BALB mice which were treated with either anti-PD-1 or control antibody after the solid tumours were established to ∼100 mm^3^ (Fig. 6d). We observed that the tumours targeted by two independent *Stub1* sgRNA were consistently more aggressive than the control tumours in mice treated with control antibody (Fig. 6e and Supplementary Fig. 8i). However, unlike the control tumours, mice bearing the *Stub1*-null tumours responded to the anti-PD-1 treatment (Supplementary Fig. 8m). But no significant differences were seen in terms of tumour volume at day 10 among the tumour-bearing mice treated with anti-PD-1 antibody (Fig. 6e). Indeed, inactivation of *Stub1* in CT26 provided no benefits towards the survival of mice (Fig. 6f).

**Fig. 6.**
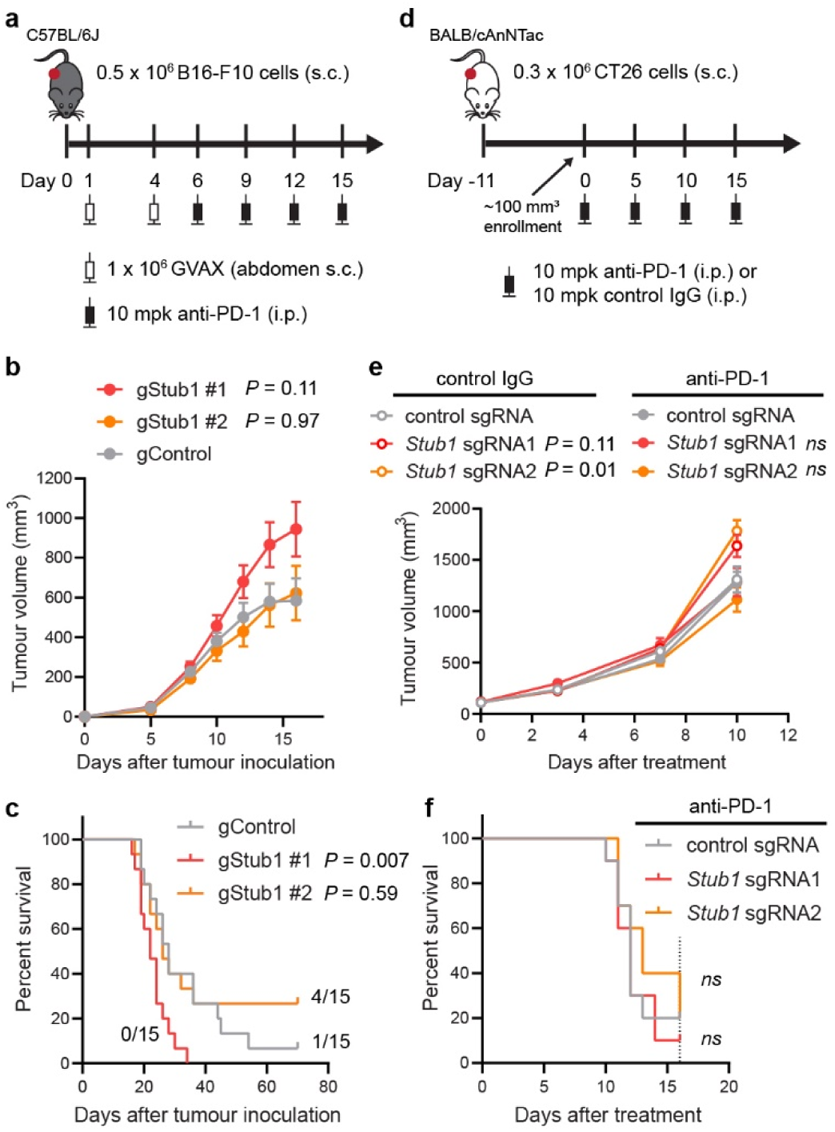
The effect of *Stub1* deletion in mouse syngeneic tumour models. **a**, Study design for C57BL mice implanted with CRISPR-edited B16-F10 clonal cells. s.c., subcutaneous. i.p., intraperitoneal. mpk, milligram per kilogram. **b**, Plot showing tumour volume of the implanted CRISPR-edited B16-F10 clonal cells. Data are mean ± s.e.m., *n* = 15 mice per group. See Supplementary Fig. 8b–d for individual tumour volume across 70 days. **c**, Kaplan-Meier survival curves of tumour-bearing mice. Median survival: gControl, 28 days; gStub1 #1, 22 days; gStub1 #2, 26 days. Either 1 or 4 out of 15 mice bearing gControl or gStub1 #2 tumour cells respectively were still alive at day 70. **d**, Study design for BALB mice implanted with CRISPR-edited CT26 cells. See Supplementary Fig. 9 for full characterization of the cells. **e**, Plot showing tumour volume of the implanted CRISPR-edited CT26 cells. Data are mean ± s.e.m. See Supplementary Fig. 8f–h and 8j–l for individual tumour volume across 16 days. **f**, Kaplan-Meier survival curves of tumour-bearing mice treated with anti-PD-1 antibody. Median survival: control sgRNA, 12 days; *Stub1* sgRNA1, 12 days; *Stub1* sgRNA2, 13 days. The study was terminated at day 16 (dotted line). See Supplementary Fig. 8i for the survival curves of mice treated with control antibody. *n* = 10 mice per group (e, f). *P* values were determined by ordinary one-way ANOVA on day 16 data (b) or two-way ANOVA on day 10 data (e) versus control tumours with Sidak’s multiple comparisons test, ns *P* ≥0.50. *P* values were determined by Log-rank (Mantel-Cox) test versus control tumours (c, f), ns *P* ≥0.50.

## Discussion

Emerging evidence^9,10,32,33,50^ points to a role for STUB1 in tumour immune evasion and anti-PD-1 resistance. However, the underlying mechanisms have been largely unclear. STUB1 protein^51^ is evolutionarily conserved among many species^52^. The protein is highly homologous (>97% identical) between human and mouse, with absolute identity at the substrate binding pocket.

Here, using several murine and human cell lines, we provided multiple lines of evidence that STUB1 downregulates IFNGR1 to dampen IFNγ sensing. During preparation of this manuscript, γ Peeper and co-workers arrived at a similar conclusion where they elegantly identified STUB1 as a pivotal regulator of IFNGR1 through CRISPR screen and further pinpointed the ubiquitination site on IFNGR1 with MS proteomics^53^. This independent finding further strengthens the case of STUB1 as an intracellular checkpoint and barrier for IFNγ sensing.

Throughout our *in vitro* experiments, STUB1 consistently constrains the IFNγ sensing across several murine and human tumour cell lines. Deletion of *STUB1* in human prostate and breast tumour cells further sensitized them to growth inhibition induced by IFNγ (Fig. 5d). ICB resistance can be conferred through defective^4,5,22^ IFNγ signaling or pathway insensitivity^4,20,23–25^. Thus, we hypothesized that inactivation of STUB1 may reverse ICB resistance by increasing tumour cells’ sensitivity for IFNγ. Nonetheless, these results did not translate to the *in vivo* murine models. In B16-F10 model, we observed conflicting survival results (Fig. 6c) between the cells targeted with two independent CRISPR guide RNAs. We attributed these to clonal effects resulting from single-cell subcloning (Fig. 2g, Supplementary Fig. 2e–f). To explore things more broadly, we also investigated whether *Stub1* deletion could reverse ICB resistance in the CT26 model. In this case, we ensured the clonal diversity was preserved before tumour inoculation. However, we did not observe significant regression of *Stub1*-null tumours relative to the control tumours or improved median survival in the murine model (Fig. 6e–f). The disconnection between the *in vitro* and *in vivo* results is multifactorial. First, we generated all the knockout lines by electroporating the gRNA/Cas9 RNP directly into cells. This non-viral approach leaves no traces behind and does not permanently introduce foreign elements, such as Cas9 protein, antibiotic resistance marker or fluorescent protein, which can artificially enhance the immunogenicity of the cells expressing them^39^. In addition, IFNγ–a pleiotropic cytokine–is a double-edged sword that not only increases the immunogenicity of the tumours and primes them for growth inhibition, but also upregulates several inhibitory immune proteins^54,55^. Finally, our understanding of STUB1, in terms of its true biological function, is still fragmented. Other biological pathways regulated by STUB1 might oppose and negate the anti-tumour immunity in a complex tumour microenvironment^56–58^.

In summary, our results highlight STUB1 as an intracellular checkpoint for IFNγ sensing. Inactivation of STUB1 increased tumour cells’ sensitivity for IFNγ, which in turn upregulated ISGs expression and enhanced antigen processing and presentation *in vitro*. We attributed these observations to the physiological role of STUB1 to downregulate the level of IFNGR1 on tumour cells’ surface, thereby reducing their ability to sense IFNγ – a key cytokine secreted by activated T cells and NK cells. Importantly, we showed that pharmacological inhibition of STUB1 with ectopic expression of a biologic phenocopied the genetic knockout, suggesting a way to target STUB1 with chemical inhibitors. Finally, upon exposure to a combination of IFNγ and TNFα, loss of STUB1 further sensitized tumour cells to growth inhibition *in vitro*. Although targeting STUB1 may offer a rational approach to improve the anti-tumour immunity when combined with anti-PD-1 treatment, we did not observe reversal of ICB resistance *in vivo*, suggesting that this type or level of pathway upregulation may not be sufficient to confer therapeutic benefit, at least in the mouse models we used. However, it should be noted that our investigations were limited to probing the role of STUB1 in syngeneic tumour cells. There may be additional anti-tumour benefit to also inhibiting STUB1 in immune cells, e.g., CD8^+^ T cell^59^, as well as the most relevant human cancer types, such as prostate and breast cancer, as implicated by the analysis of the TCGA dataset. As such, a specific chemical tool that could systemically interrogate the role of STUB1 in the peripheral immune system and tumour microenvironment is highly desirable and would complement the genetic approach. These approaches would be central to the further exploration of STUB1 as a potential immuno-oncology target.

## Acknowledgements

We thank MSD for the financial support. SN acknowledges support from the MRL Postdoctoral Research Program. We thank Byung Lee and Chunhong Shao from HD Biosciences for executing animal studies.

## Methods

### Protein and peptides

Recombinant STUB1 protein, spanning aa25-aa153, was produced by Nanyang Technological University protein production platform. The purity and identity of the protein was confirmed by mass spectrometry and SDS-PAGE. Synthetic peptides, in a form of N-acetylation and free C-terminal carboxylic acid, were custom made by Chinese Peptide Company (CPC). The purity and identity of the peptides were confirmed by analytic HPLC (≥95% purity) and mass ≥ spectrometry. Peptides are dissolved in neat DMSO as 10 mM stock solution and diluted thereof for subsequent experiments.

### Cell lines and culture

Murine melanoma B16-F10 (CRL-6475), murine colon CT26 (CRL-2638), human melanoma A375 (CRL-1619), human lung A549 (CCL-185), human prostate DU145 (HTB-81), human prostate PC-3 (CRL-1435) and human breast MCF7 (HTB-22) were purchased from ATCC. B16-F10 or A375 were cultured in DMEM (Gibco, #10569010). CT26 was cultured in RPMI-1640 (Gibco, # 72400047). A549 and PC-3 cells were cultured in Ham’s F-12K (Gibco, #21127022). DU145 and MCF7 were cultured in MEM (Gibco, # 2360099). All culture media are supplemented with 10% FBS (HyClone, #SH30071.03). Human recombinant insulin (10 μg ml^−1^, Gibco #12585014) was additionally included in the culture media for MCF7. All cells were maintained at 37 °C, 5% CO_2_, and 95% relative humidity. The cells were routinely tested for mycoplasma. The CRISPR-edited B16-F10 and CT26 cell lines were PCR-evaluated by IDEXX BioAnalytics to be free of viral contamination. The CRISPR-edited B16-F10 lines were genetically confirmed as mouse origin, and had almost identical short tandem repeat profile (>90% match) to that established for B16-F10 (ATCC, CRL-6475). Cell number was determined using NC-100 NucleoCounter (ChemoMetec).

### Virus-free generation of gene-knockout cell lines

Tumour cells were genetically edited by electroporating the Cas9/crRNA/tracrRNA or Cas9/sgRNA ribonucleoprotein (RNP) complexes into cells using the 4D-nucleofector system (Lonza). To prepare the guide RNA complex, a 1:1 mixture of Alt-R® crRNA and tracrRNA (50 μM each, IDT) in nuclease-free duplex buffer (IDT) was heated at 95 °C for 5 min, followed by cooling to room temperature. The crRNA/tracrRNA complex or the sgRNA (150 pmol) was mixed with Alt-R® S.p. HiFi Cas9 Nuclease V3 (100 pmol, IDT, #1081060), and the resulting mixture was incubated at room temperature for 10 min to form the final RNP complexes which were used immediately. In parallel, the harvested tumour cells were rinsed with PBS. After re-suspending in 20 μl nucleofactor solution (Lonza), the cell suspension was added to the final RNP complexes (4.6 μl) in microcentrifuge tube. The resulting cell suspension was transferred to a designated well of nucleocuvette strip which was then pulsed with the nucleofector system using a preset program. After pulsing, culture media (75 μl) was added and the cell suspension was transferred to a designated well of a 12-well plate filled with 1.0 ml culture media. After 48 h incubation, the CRISPR-edited B16-F10 were subcloned by limiting dilution. The monoclonal cell lines were validated by Western blotting and analyzing the Sanger sequencing results of the PCR amplicon (∼800 bp) flanking the crRNA-targeted site using ICE v2 CRISPR analysis tool (Synthego). For CT26, the cells were sorted for the top 50^th^ percentile of IFNGR1-high population using BD FACSAria to enrich for the *Stub1*-null cells (Supplementary Fig. 9a).

Human tumour cells, e.g., DU145, PC-3 and MCF7, were used right after the CRISPR editing. Sorting or subcloning was not applied to the human cell lines, since these lines were studied for a brief period in *in vitro* setting. Loss of STUB1 protein in the human cell lines was confirmed by Western blot analysis.

**Parameter and conditon of eletroporation:**

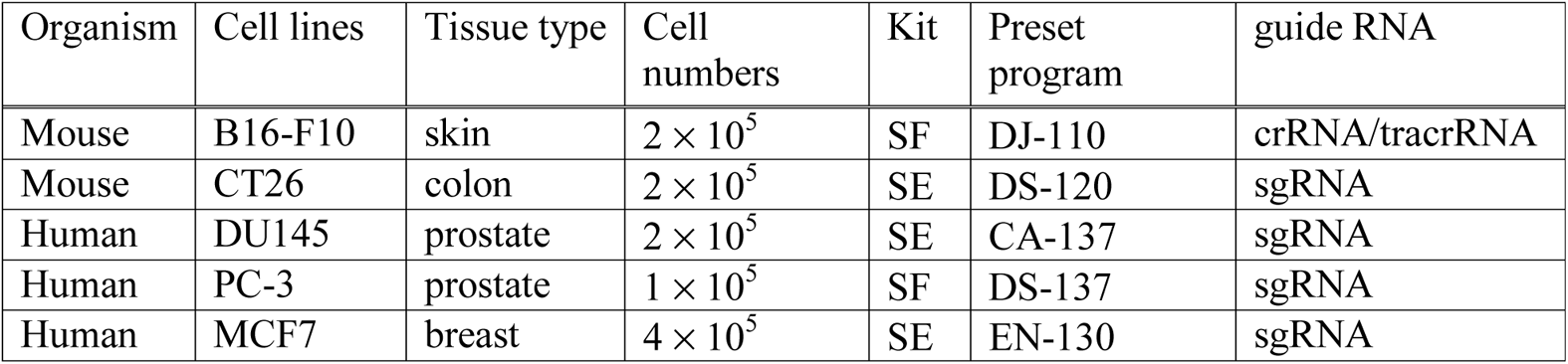

*Stub1* crRNA1 or sgRNA1: GCATTGCTAAGAAGAAGCGC;

*Stub1* crRNA2 or sgRNA2: ACTTGCGGCCCACGAAGAGC;

control crRNA or sgRNA: GCGAGGTATTCGGCTCCGCG;

*STUB1* sgRNA: GGCCGTGTATTACACCAACC;

*PTPN2* sgRNA: CCACTCTATGAGGATAGTCA.

### Generation of tumour cell lines expressing mCherry2-peptide fusion

To generate the plasmids, gBlocks® gene fragments (IDT) encoding the inhibitory biologic (FLAG-mCherry2-GGSGGS-SIWWPD) and the control biologic (FLAG-mCherry2-GGSGGS-SIWWHR) were cloned into pEF6 vector (Thermo Fisher Scientific) by standard restriction enzyme digestion and T4 DNA ligation. The final constructs were verified by Sanger sequencing. Coding sequences of the constructs are provided in Supplementary Table 5. To generate the stable cell lines, a total of 2 × 10^5^ B16-F10, A375 or A549 cells were electroporated with 200 ng plasmid using DJ-110, FF-120 or CM-130 respectively – the preset programmed in 4D-nucleofector system (Lonza). The preparation of the cell suspension and the process of the electroporation are similar to that described in the CRISPR method section. The stable cell lines were selected using 10 μg ml^−1^ blasticidin three days post-electroporation and were maintained in 5 μg ml^−1^ blasticidin once the stable colonies were established.

### In vitro stimulation with IFN**γ**

Parental, CRISPR-edited or biologic-overexpressed tumour cells were seeded in a cell density of 60,000 (B16-F10), 150,000 (CT26), 100,000 (A375), or 200,000 (A549) per well in 12-well plate filled with 0.8 ml culture media + 10% FBS. After overnight incubation, the culture media were replaced with 1 ml fresh media supplemented with 10% FBS and the designated concentration of the recombinant mouse IFNγ (R&D Systems, 485-MI-100) or recombinant human IFNγ (R&D Systems, 285-IF-100). The cells were stimulated with the cytokine for 24 h before they were harvested by trypsinization for flow cytometry, western blot, or qPCR analysis (6 h treatment).

### Flow cytometry analysis

Cells were dissociated from the wells with 0.25% trypsin (Gibco, 25200056). After rinsing with 0.5 ml PBS, each sample was stained with the LIVE/DEAD Fixable Aqua dead cell staining solution (Life Technologies, L34957) in 100 μl PBS (1:1000 dilution) for 15 min at 4 °C. After rinsing with 2 × 0.5 ml PBS, each sample was stained with the corresponding primary antibody-dye conjugates diluted in 100 μl Pharmingen stain buffer (BD Biosciences, #554657). After 1 h staining at 4 °C, the cells were rinsed with 2 × 2 ml Pharmingen stain buffer and fixed with 4.21% (w/w) formaldehyde (BD Biosciences, #554655). Cells were analyzed on LSRFortessa X-20 (BD Biosciences) with appropiate fluorescence compensation. The data were analyzed using FlowJo. Cells were gated based on FSC and SSC. Single cells were selected using FSC-A and FSC-H and viable cells were selected using LIVE/DEAD signal. Primary antibodies used were: H-2K^b^/H-2D^b^ (FITC, Biolegend, #114606, 1:50 dilution), HLA-A,B,C (FITC, Biolegend, #311404, 1:100 dilution), mouse IFNGR1 (PE, Invitrogen, #12-1191-82, 1:50 dilution), human IFNGR1 (PE, Miltenyi, #130-125-851, 1:50 dilution), mouse IL1R1 (APC, Biolegend, #113509, 1:20 dilution), mouse IL6R (APC, Biolegend, #115812, 1:20 dilution), mouse IFNAR1 (APC, Biolegend, #127314, 1:20 dilution), and mouse GP130 (PE, Biolegend, #149404, 1:100 dilution).

### Western blot analysis

Cells were dissociated from the wells with 0.25% trypsin (Gibco, 25200056). After rinsing with 0.5 ml PBS, the cell pellets were lysed with chilled cell lysis buffer (Cell Signaling Technology, #9803) supplemented with Halt− protease inhibitor cocktail (Thermo Scientific, #78430) and phosphatase inhibitor (Sigma-Aldrich, # 4906837001) for 30 min with intermittent vortexing. The lysate was transferred into PCR-strip tubes and sonicated in a chilled water bath sonicator (QSonica). Lysates were clarified by centrifugation at 15,000 rpm at 4 °C for 15 min. Protein concentration was determined using BCA protein assay kit (Pierce, #23225). The lysates were mixed with LDS sample buffer (Life Technologies, NP0008) and sample reducing agent (Life Technologies, NP0009), followed by heating at 70 °C for 10 min to fully denature the protein. The protein extract (20 μg) was separated on 4–12% NuPAGE Bis-Tris gels (Life Technologies WG1403A), followed by transferring onto nitrocellulose membranes using the Trans-Blot® Turbo™ semi-dry system (Bio-Rad). Membrane blots were pre-stained with total protein stain (LI-COR, # 926-11016) and imaged with Odyssey® CLx. The blots were subsequently blocked for 1 h at room temperature with Intercept® (TBS) blocking buffer (LI-COR, # 927-60001). The blots were finally probed, for overnight at 4 °C, with the appropriate primary antibodies diluted in Intercept® (TBS) blocking buffer supplemented with 0.1% (v/v) Tween-20, followed by the secondary antibodies (IRDye® 800CW donkey anti-rabbit or anti-mouse IgG, LI-COR) for 1 h at room temperature. Fluorescent signals were imaged and quantified using Odyssey® CLx and Image Studio v5.0. Primary antibodies used were: STUB1/CHIP (Cell Signaling Technology, #2080, 1:2000 dilution), STAT1 (Cell Signaling Technology, #14995, 1:10,000 dilution), phospho-Tyr701-STAT1 (Cell Signaling Technology, #9167, 1:2,000 dilution), STAT2 (Cell Signaling Technology, #72604, 1:2000 dilution), IRF1 (Cell Signaling Technology, #8478, 1:2000 dilution), PSMB8 (Cell Signaling Technology, #13635, 1:2,000 dilution), PSMB9 (Abcam, ab184172, 1:10,000 dilution), and PSMB10 (Abcam, ab183506, 1:10,000 dilution).

### qPCR analysis

Total RNA was extracted from the tumour cells using RNeasy plus mini kit (Qiagen, #74134) according to the manufacturer’s instructions. A total of 1 μg RNA was reversely transcribed in a 20 μl reaction mixture using high-capacity cDNA reverse transcription kit (Applied Biosystems, #4368814) according to the manufacturer’s instructions. The resulting cDNA mixture was diluted to 100 μl with nuclease-free water and an aliquot of 2 μl was used for each qPCR set-up. The qPCR was conducted with QuantStudio 12K Flex using power SYBR™ green PCR master mix (Applied Biosystems, #4368577) and 500 nM primer set (IDT PrimeTime) in a total volume of 10 μl reaction in 384-well plates. The PCR cycle is as follow: incubation at 95 °C (10 min), followed by 40 cycles of 95 °C (15 sec) and 60 °C (60 sec). Four technical replicates were performed in parallel for each biological replicate. ΔCT was calculated by taking the difference between the mean CT value (*n* = 4) for each gene of interest and the mean CT value (*n* = 4) of a reference gene (gene name: *Tbp*) within a biological sample. Fold change in gene expression wa derived from the ÄÄCT using untreated gControl cells as the reference. PCR primer set for *Ifngr1*: ATGATCAGAAATGTTGGTGCAG and TTGAACCCTGTCGTATGCTG; *Stat1*: GACTTCAGACACAGAAATCAACTC and TTGACAAAGACCACGCCTT; *Irf1*: ACTCAGACTGTTCAAAGAGCTTC and GTCACCCATGCCTTCCAC; *Tbp*: CCAGAACTGAAAATCAACGCAG and TGTATCTACCGTGAATCTTGGC.

### Gene expression profilling with NanoString

CRISPR-engineered B16-F10 cells (gControl, gStub1 #1, and gStub1 #2) were seeded separately in 12-well plate (50,000 cells per well) with DMEM + 10% FBS. After overnight incubation, the culture media were replaced with fresh media (DMEM + 10% FBS) supplemented with 0.03 ng ml^−1^ of recombinant mouse IFNγ (R&D Systems, #485-MI-100). Total RNA from untreated cells (24 h) and IFNγ-treated cells (6 or 24 h) were extracted with RNeasy plus mini kit (Qiagen, 74134) according to manufacturer’s protocol. An input of 150 ng RNA from each sample was mixed with the NanoString reporter and capture probes (nCounter Mouse PanCancer IO 360, # XT-CSPS-MIO360-12), and incubated at 65 °C for 20 h. The hybridized samples were processed on the nCounter prep station, and the resulting cartridge was scanned by the nCounter digital analyzer using 555 fields of view. Raw count data were evaluated for quality control and normalized with 19 housekeeping genes using nSolver 4.0 software (Supplementary Table 3).

*Tlk2* was excluded from the housekeeping gene due to weak expression (RNA counts <80). Fold change (FC) was calculated by comparing the normalized RNA counts of each sample to that of the untreated gControl cells as the denominator (Supplementary Table 3). Weakly expressed genes, where the normalized RNA counts were consistently less than 80 in all samples, were excluded from fold change analysis, resulting in an evaluable set of 493 out of 750 genes (Supplementary Fig. 4a). As an overview, all 750 targeted genes were included in the scatter plot analysis (Supplementary Fig. 4b–d).

### Proteomics by mass spectrometry

CRISPR-engineered B16-F10 cells (gControl, gStub1 #1, and gStub1 #2) were seeded separately in 12-well plate (50,000 cells per well) with DMEM + 10% FBS. After overnight incubation, the culture media were replaced with fresh media (DMEM + 10% FBS) supplemented with 0.03 ng ml^−1^ of recombinant mouse IFNγ (R&D Systems, #485-MI-100). After 24 h treatment, the cells were trypsinized, collected and washed twice with 0.5 ml PBS. The cell pellets collected from three independent experiments on separate day were lysed in 100 μl of lysis buffer containing 4% sodium dodecyl sulfate, 50 mM Tris-HCl pH 7.5, 50 μg ml^−1^ of DNase (Roche, #10104159001), 50 μg ml^−1^ of RNase (Roche, #10109169001) and Halt− protease inhibitor cocktail (Thermo Scientific, #78430) on ice for 30 min followed by 65 °C for 30 min. Lysates were clarified by centrifugation at 16,000g at 10 °C for 30 min. Protein content was re-extracted from the pellet with 50 μl of lysis buffer, sonicated with a single burst using a probe sonicator and heated at 95 °C for 10 min before centrifugation at 16,000g at 10 °C for 15 min. Lysates from first and second extractions were pooled. Protein concentration was determined using BCA protein assay kit (Pierce, # 23225). Detergent removal and protein digestion were performed in centrifugal suspension trap columns (Protifi, C02-micro) according to the manufacturer’s instructions. Briefly, 80 μg protein extract from each sample was reduced with 50 mM dithiothreitol (Sigma, #43815) at 95 °C for 10 min and then alkylated with 100 mM iodoacetamide (Sigma, I2512) in the dark at ambient temperature for 30 min. The samples were acidified with 1.2% of phosphoric acid and mixed well with washing buffer consisting of 90% methanol and 100 mM ammonium bicarbonate. The samples were transferred to the suspension trap columns and centrifuged at 4000 g for 30 s. The columns were washed 3 times with washing buffer. Proteins trapped in the suspension bed were digested in 50 mM ammonium bicarbonate with trypsin and endoproteinase Lys-C (Promega, V5073) at enzyme to protein ratio 1:25 in a 47 °C waterbath for 2 h. Peptides were eluted firstly with 50 mM ammonium bicarbonate, then with 0.2% formic acid and lastly with 50% acetonitrile and 0.2% formic acid. The eluates were pooled and vacuum dried completely. Dried peptides were reconstituted with 0.1% formic acid in water, followed by injecting 4 μg for mass spectrometry analysis. Peptides were loaded on a reverse phase EASY-Spray− column (50 cm × 75 μm inner diameter) operated using Easy-nLC− 1200 μ (Thermo Fisher Scientific) coupled online to a Q-Exactive HF-X mass spectrometer (Thermo Fisher Scientific). Peptides were separated using a 120 min linear gradient at a flow rate of 300 nL min^−1^. The Q-Exactive was operated in ‘top-10’ data-dependent acquisition (DDA) mode with full scan acquired at a resolution of 120,000 (scan range 200–1800 m/z) with an automatic gain control (AGC) target of 3e6. The top ten most abundant ions from the full scan were isolated with an isolation width of 0.7 m/z and fragmented by higher energy collisional dissociation (HCD) with normalized collision energy (NCE) of 27. MS/MS scan was acquired at a resolution of 30,000 with an AGC target of 1e5. The default charge state was set at 2 and dynamic exclusion was enabled for 10 s. Maximum ion injection time for full scan and MS/MS scan were 100 ms and 105 ms respectively.

DDA raw files were processed with Proteome Discoverer 2.4 using Sequest HT search engine where mass spectrometric data was searched against SwissProt TaxID 10090 mouse database (v2017-10-25). Percolator was used to validate search results based on the concatenated mode where only the best scoring PSMs (target/decoy) were considered. Trypsin was specified as the enzyme, cleaving after all lysine and arginine residues and allowing up to two missed cleavages. Carbamidomethylation of cysteines was set as fixed modification while variable modifications included oxidation of methionine, acetylation of N-terminus, N-terminal loss of methionine and N-terminal loss of metionine along with the addition of an acetyl group. The minimum peptide length required for protein identification was six amino acids. Precursor and fragment mass tolerances were set as 10 ppm and 0.02 Da respectively. Overall, a total of 3048 proteins were detected by mass spectrometry (*n* = 6 replicates per cell group, 3 biological replicates × 2 mass spectrometry replicates). After quality control (>1 unique peptide found or 1 unique peptide with ≥ 25% coverage), we obtained a high-quality dataset of 2293 proteins for further differential enrichment analysis (Supplementary Table 4). The adjusted *P* values were determined by unpaired t test per protein (without assuming a consistent standard deviation) and false discovery rate approach (two-stage step-up method of Benjamini, Krieger, and Yekutieli, with Q = 5%).

Differentially expressed proteins are defined by Log_2_ (Fold change) >1 and −Log_10_ (adjusted *P*) >1.301 (Supplementary Table 4).

### MHC-I peptide immunoprecipitation

B16-F10 cells were cultured in DMEM supplemented with 10% FBS in 2-500 cm^2^ triple layer flasks for 24 hours in 0.03 ng ml^−1^ recombinant mouse IFNγ. At the time of harvest, cells were washed with PBS and then lifted using PBS-based Enzyme Free Cell Dissociation Solution (Sigma-Aldrich). Cells were pelleted at 500 x g for 5 minutes and then resuspended in 4 mL lysis buffer [25 mM Tris-HCl pH 7.4, 150 mM NaCl, 1 mM EDTA, 1% NP-40, 5% glycerol) +1x HALT protease/phosphatase inhibitors (Pierce)] and equilibrated on ice for 30 minutes. The supernatant was quantified for protein concentration using the BCA Assay (Pierce) and 30 mg protein lysate was used for the IP.

Immunoprecipitations were performed using an automated liquid handler (AssayMAP Bravo, Agilent Technologies) as described in *Mol. Cell. Proteom.* (2021) 20, 100108. Briefly, 0.25 mg anti-H2^Db^ (B22.249, Thermo Fisher Scientific) and 0.25 mg anti-H2^Kb^ (Y3, BioXCell) were immobilized on each Protein-A cartridge (25 µL bed volume, Agilent Technologies, cat. no. G5496-60018) and crosslinked. Each lysate sample was divided and loaded onto two of these cartridges at 20 µL/min before washes with TBS supplemented with 0.2 M NaCl and 25 mM Tris-HCl (pH 7.4) and then eluted with 1% acetic acid. Eluates were desalted using C18 cartridges (5 µL bed volume, Agilent Technologies, cat. no. 5190–6532) on an automated liquid handler as per the manufacturer’s instructions. Peptides were dried and stored at −80°C until analysis.

### Analysis of MHC-bound peptides by mass spectrometry

Dried peptides were dissolved in 3% acetonitrile (ACN) with 0.1% formic acid (FA), injected onto an EASY-Spray analytical column (C18, 75_μm i.d. × 50 cm, 2 μm particle size; Thermo Fisher Scientific) at a flow rate of 0.3 ng/µL and separated chromatographically using a linear gradient of 3-45% B (A= 0.1% FA, B= 99.9% ACN, 0.1% FA) in 120 minutes. Mass spectra were detected using an Orbitrap Fusion Lumos in data-dependent acquisition mode at resolution of 60,000 with an AGC target of 1E6. MS/MS spectra were acquired in both HCD and CID mode with collision energies of 30% and 35%, respectively, with an AGC of 1E4 and maximum injection times of 100 ms and resolution of 7,500.

Raw mass spectral files were analyzed using MaxQuant v1.6.1.0 and searched against the mouse SwissProt reference database (Proteome ID: UP000000589, 55,366 entries; downloaded June 7, 2020). Methionine oxidation and protein N-terminal acetylation were set to variable modifications and the digestion mode was set to unspecific. The matching-between-runs option (0.4 min match time window) was enabled. Search results were filtered for peptides 8-12 amino acids in length and 5% peptide FDR. Statistical data analysis and filtering was performed using Perseus software v1.6.15.0 and plotted using GraphPad Prism 8.

### Isothermal titration calorimetry (ITC)

Recombinant STUB1 protein (aa25-aa153) was dialyzed overnight with Slide-A-Lyzer cassette (7K MWCO, Thermo Scientific, #66373) in 1 liter of dialysis buffer (PBS, pH 7.4, 0.5 mM TCEP). The dialyzed protein solution was centrifuged at 15,000 rpm for 10 min at 4 °C to remove potential precipiates. The protein was diluted to 20 μM using the dialysis buffer, followed by the additon of DMSO spike-in (2% final concentration). The synthetic peptides (10 mM in DMSO) were diluted to 200 μM with the dialysis buffer (2% final DMSO concentration). ITC measurements were performed at 25 °C using a Microcal PEAQ-ITC (Malvern Panalytical Inc). An initial injection of 0.4 µl followed by a total of 39 injections of peptide solution (1 µl, 200 μM) were added at an intervals of 2 min into the protein solution (20 μM) while stirring at 750 rpm. The data point produced by the first injection was discarded prior to curve fitting in order to account for the diffusion effect during the equilibration process. The experimental data were fitted to a non-interacting one-site binding model using the analysis software supplied by Microcal, with ΔH (enthalpy change), K_a_ (association constant) and N (number of binding sites per monomer) as adjustable parameters. Free energy change (ΔG) and entropy contributions (TΔS) were determined from the standard equation: ΔG = ΔH–TΔS = –RT lnK_a_, where T is the absolute temperature and R = 1.987 cal mol^−1^ K^−1^.

### Thermal shift assay

The SYPRO Orange fluorescent dye (Invitrogen) was used to measure the thermal stability of recombinant STUB1 protein (aa25-aa153). With increasing temperature, binding of the dye molecule to the hydrophobic region of the denatured STUB1 results in an increase in the fluorescence intensity. The midpoint of this transition is termed the T_m_. Binding of a ligand, such as peptide, stabilizes the protein and results in a melting temperature shift (⊗T_m_), which correlates with the binding affinity of the ligand. The thermal shift assay was conducted in a CFX96™ real-time PCR detection system (Bio-Rad). A total of 50 µl mixture containing 3.125× SYPRO Orange (Invitrogen, diluted from 5000× DMSO stock), 100 μM peptide of interest, and 10 µM protein was prepared in a PCR 8-well strip tube. The samples were heated from 25 to 95 °C in 0.5 °C increment each cycle. The holding time for each cycle is 5 sec, after which the fluorescence intensity was measured in Channel 2 (HEX) with Ex/Em:515–535/560–580 nm. Each independent experiment was performed in technical duplicates.

### Competitive fluorescence polarization

The assays were performed at room temperature using assay buffer (PBS, pH 7.4, 0.01% v/v Tween 20) and black 384-well non-binding polystyrene microplate (Greiner Bio-one, #784900). The peptide of interest was first diluted (10-point, 3-fold serial dilution) with the assay buffer on the microplate to have a volume of 10 µl in each well. This was followed by the addition of 10 µ mixture containing 5-FAM-SSGPTIEEVD-CO_2_H (30 nM) and the recombinant STUB1 protein (2 μM). The final assay solution (20 µl) contains 5-FAM-labeled tracer peptide (15 nM), protein (1 μM) and peptide of interest (5 nM to 100 µM). After 30 min incubation in the dark, the microplate was read with TECAN Infinite M1000 PRO (Ex: 470 nm, Em: 520 nm, bandwidth: 5 nm, G-factor = 1.05, gain: optimal, #flashes = 10, settle time = 0 ms, z position: calculated from well). Value of polarization (mP) = 1000 × (G × intensity – intensity⊥) / (G × intensity + intensity⊥). Half maximal inhibitory concentration (IC_50_) was determined by fitting the curve using 4-parameter sigmoidal function in GraphPad Prism. Each independent experiment was performed in technical triplicates.

### Co-immunoprecipitation of FLAG-mCherry2-peptide and STUB1

B16-F10 cells stably expressing the biologic were harvested, rinsed with PBS, and lysed with chilled cell lysis buffer (Cell Signaling Technology, #9803) supplemented with Halt− protease inhibitor cocktail (Life Technologies, #78430) and phosphatase inhibitor (Sigma-Aldrich, # 4906837001). Cellular lysates were clarified by centrifugation at 15,000 rpm at 4 °C for 15 min.

Protein concentration was determined using BCA protein assay kit (Pierce, #23225). For each sample of the co-immunoprecipitation (co-IP), 20 μl of anti-FLAG magnetic beads (Sigma-Aldrich, M8823) was rinsed twice with 0.2 ml PBS, followed by addition of 500 μl diluted cellular lysate (60 μg, 0.12 μg μl^−1^). For competitive inhibition, synthetic peptide (SIWWPD) was added into the co-IP mixture. The resulting mixture was rotated at room temperature for 4 h, after which the beads were rinsed with 3 × 0.5 ml PBS to remove the unbound proteins. Bound protein complexes were directly eluted with a 20 μl solution of LDS sample buffer (Life Technologies, NP0008) supplemented with sample reducing agent (Life Technologies, NP0009), followed by heating at 70 °C for 10 min. The co-IP final extract was separated on 4–12% Bolt Bis-Tris gels (Life Technologies). Blotting was similar to the Western Blot section described above. Primary antibodies used were: STUB1/CHIP (Cell Signaling Technology, #2080, 1:1000 dilution), FLAG (Sigma-Aldrich, F1804, 1:1000 dilution). As a comparison, 30 μg of whole cell lysates (half amount for the input of co-IP) were loaded along with the co-IP final extract in the gel.

### The Cancer Genome Atlas dataset analysis

The Cancer Genome Atlas (TCGA) database was used for analysis of clinical relevance. RNA-sequencing data for 9963 tumors and somatic alterations data for 6384 tumors were obtained through TCGA portal (https://portal.gdc.cancer.gov/) as of September 2015. The expression data were Log10 transformed. Spearman correlation was used to determine the correlation and Wilcoxon rank-sum test was used to calculate *P* value. Statistical analyses and visualizations were performed with Matlab R2010b Version 7.11.2. TMB cutoff for the pan-tumor clinical cohort was the Youden Index value derived in AUROC analysis. An additional, exploratory, pan-tumor TMB threshold was derived by using TMB and GEP data, similar to a previously described method^60^.

### Lumit immunoassay measuring p-STAT1 level

Parental and CRISPR-edited tumour cells were seeded in a cell density of 12,500 (DU145), 15,000 (PC-3), or 30,000 (MCF7) per well in 384-well plate (Greiner Bio-One, #781080) filled with 25 μl culture media + 10% FBS. After 20–24 h incubation, the cells were treated with recombinant human IFNγ (R&D Systems, 285-IF-100) by adding an equal volume of the complete culture media containing the cytokine. After 30 min of stimulation, the culture media were removed by gentle spin using Blue Washer (BlueCatBio). Lumit immunoassay (Promega, #W1202) was performed according to manufacturer’s instrutions. The assay buffer was supplemented with Halt− protease and phosphatase inhibitor cocktail (Thermo Scientific, #78443). The cells were lyzed with 12 μl per well of lysis solution (0.02% digitonin) for 20 min. The cell lystates were probed for 90 min with 12 μl of mixture of primary antibodies, which include anti-STAT1 rabbit antibody (Cell Signaling Technology, #14994), anti-pSTAT1 (Tyr701) mouse antibody (Abcam, ab29045), Lumit anti-mouse antibody-LgBiT (Promega) and Lumit anti-rabbit antibody-SmBiT (Promega). Final concentration of each antibody is 0.15 μg ml^−1^. After 2 min incubation in the presence of the Lumit detection reagent (6 μl), the luminescence signals were detected by Infinite M1000 PRO (Tecan).

### Growth inhibition measured by CellTiter-Glo and CellTiter-Fluor assay

Parental and CRISPR-edited tumour cells were seeded in a cell density of 1,800 (DU145), 2,000 (PC-3), or 10,000 (MCF7) per well in 96-well CellCarrier plate (PelkinElmer, #6005550) filled with 50 μl culture media + 10% FBS. After 20–24 h incubation, the cells were treated with 10 ng ml^−1^ of recombinant human IFNγ (R&D Systems, 285-IF-100), 10 ng ml^−1^ of recombinant human TNFα (R&D Systems, 210-TA-020/CF) or a combination of both, by adding an equal volume of the complete culture media containing the cytokines. After incubating for 6 days without media change, the CellTiter-Glo 2.0 (Promega, G9243) or CellTiter-Fluor (Promega, G6082) assay were performed according to manufacturer’ instructions. The luminescence or fluorescence (Ex: 380–400 nm, Em: 505 nm) were detected by Inifinte M1000 PRO (Tecan). For CellTiter-Fluor, triplicate wells without cells were included to determine background fluorescence and the signal average was used for background substraction.

### Syngeneic mouse studies

#### B16-F10 model

Female C57BL/6J mice (stock no: 000664, Jackson Laboratory) between 7–8 weeks of age weighing approximately 18–22 g were used for the study. The *Stub1*-null or control B16-F10 tumour cells were inoculated subcutaneously into the right lower flank with the single cell suspension of >95% viable tumour cells (0.5 × 10^6^ cells) in 0.1 ml of serum-free and phenol-red-free DMEM. One day after tumour inoculation, mice were dosed subcutaneously into the abdomen with 1.0 × 10^6^ per 100 μl GM-CSF-secreting B16 (GVAX) cells that had been irradiated with 35Gy from a ^137^Cs source discharging 124 rads min^−1^. GVAX treatment was repeated on day 4. Subsequently, mice were treated with 10 mg kg^−1^ of anti-PD-1 antibody (muDX400) *via* intraperitoneal injections on day 6. The treatment was repeated on day 9, 12, and 15. The start of the study where tumour inoculation is conducted is designated as day 0.

#### CT26 model

Female BALB/cAnNTac mice (Taconic) between 8–10 weeks of age weighing approximately 19–22 g were used for the study. The *Stub1*-null or control CT26 tumour cells were inoculated subcutaneously into the right lower flank with the single cell suspension of >95% viable tumour cells (0.3 × 10^6^ cells) in 0.1 ml of serum-free and phenol-red-free RPMI-1640. The mice were treated with 10 mg kg^−1^ of anti-PD-1 antibody (muDX400) or isotype control antibody. Drug treatment was started for all groups when mice bearing the control tumours reached an average tumor size of approximately 100 mm^3^. The start of the treatment was designated as day 0. The treatment was repeated every 5 days, for a total of 5 doses.

All animals were weighed and assigned to treatment groups using a randomization procedure. Each group has approximately the same mean animal weight. Any B16-F10 or CT26 tumours that completely or partially grow intradermally or intramuscularly were not used for the study. Irregularly shaped (e.g., W-or U-shaped) tumours were also not used. Tumours were measured in two dimensions using a caliper, and the volume was expressed in mm^3^ using the formula: V = 0.5 (*a* × *b*^2^) where *a* and *b* are the long and the short diameters of the tumor, respectively. Body weights were taken twice per week. The body weight of all mice was not significantly changed by the end of the study. Mice in a continuing deteriorating condition or with a tumour exceeding 2000 mm^3^ were considered endpoints at which the mice were euthanized by CO_2_ inhalation followed by cervical dislocation. All experiments in mice were conducted in accordance with regulations of the AAALAC and were approved by the IACUC.

### Statistical analysis

Except for the public RNAseq data, all statistical analyses were performed using GraphPad Prism 8.1.1 (GraphPad) and were described in the Figure caption.

---

**Supplementary Fig. 1.**
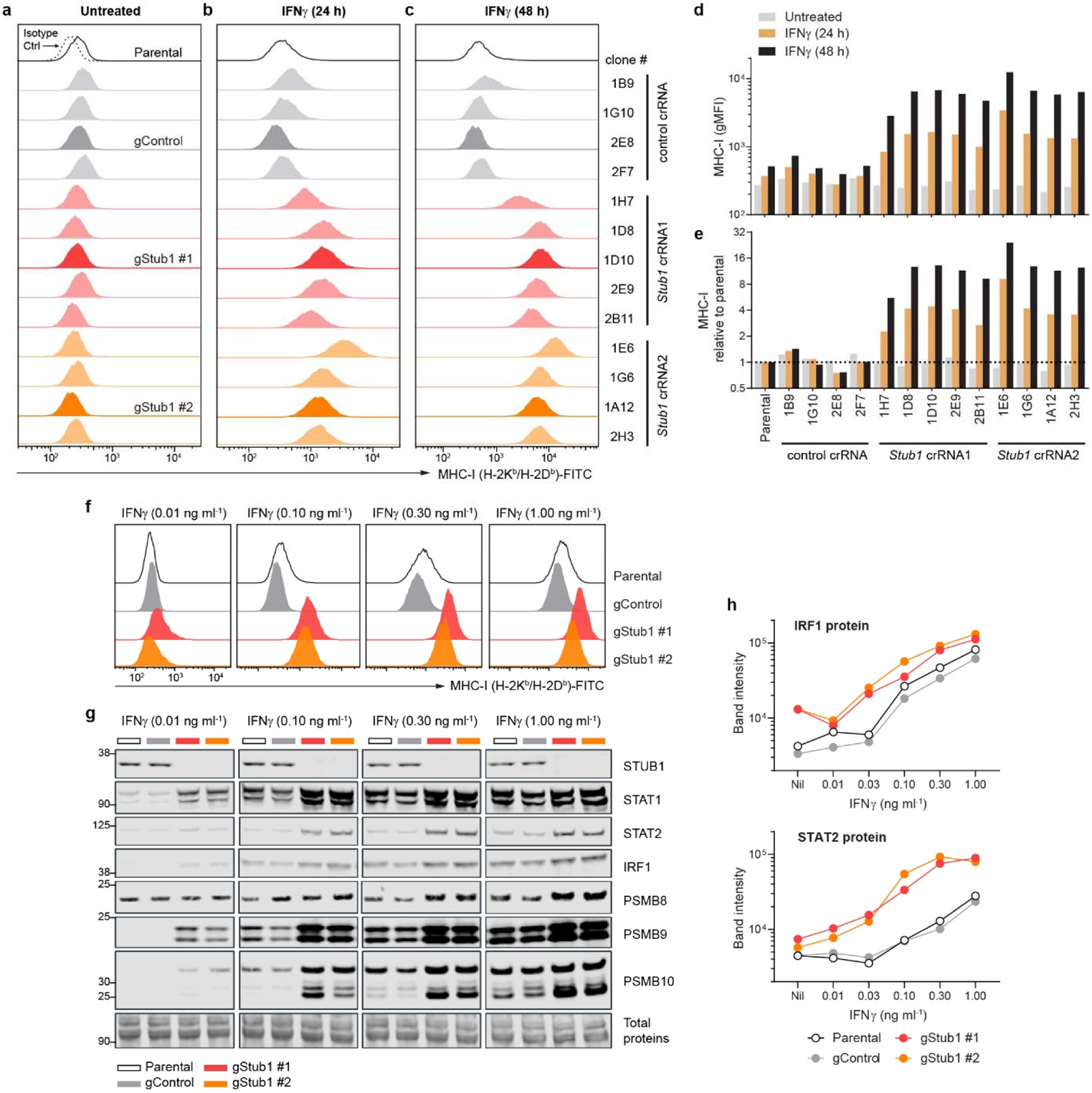
Loss of *Stub1* in B16-F10 melanoma increased the surface level of MHC-I and the protein level of STAT1, STAT2, IRF1, PSMB8, PSMB9 and PSMB10 in response to IFNγ. Related to Fig. 1. a–e, Flow cytometry analysis of cell surface level of MHC-I on parental B16-F10 and all CRISPR-edited clones isolated by single-cell subcloning (Supplementary Table 2). The tumour cells were either untreated (a) or treated with 0.10 ng ml^−1^ IFNγ for 24 h (b) or 48 h (c). The expression level of MHC-I on the tumour cells (d) and their relative abundance compared to the parental B16-F10 cells (e). All further experiments were performed using single-cell clone 2E8, 1D10 and 1A12 – termed gControl, gStub1 #1 and gStub1 #2 respectively. f, Flow cytometry analysis of cell surface MHC-I on parental, control or independent *Stub1*-null B16-F10 cells, following treatment with the indicated condition for 24 h. g, h, Western blot analysis of STUB1, STAT1, STAT2, IRF1, PSMB8, PSMB9 and PSMB10 in tumour cells, following treatment with the indicated concentration of IFNγ for 24 h (g). Quantification of the protein level with LI-COR Image Studio (h). Band intensity was normalized with total protein signal. Representative of four (f) or two (g, h) independent experiments.

**Supplementary Fig. 2.**
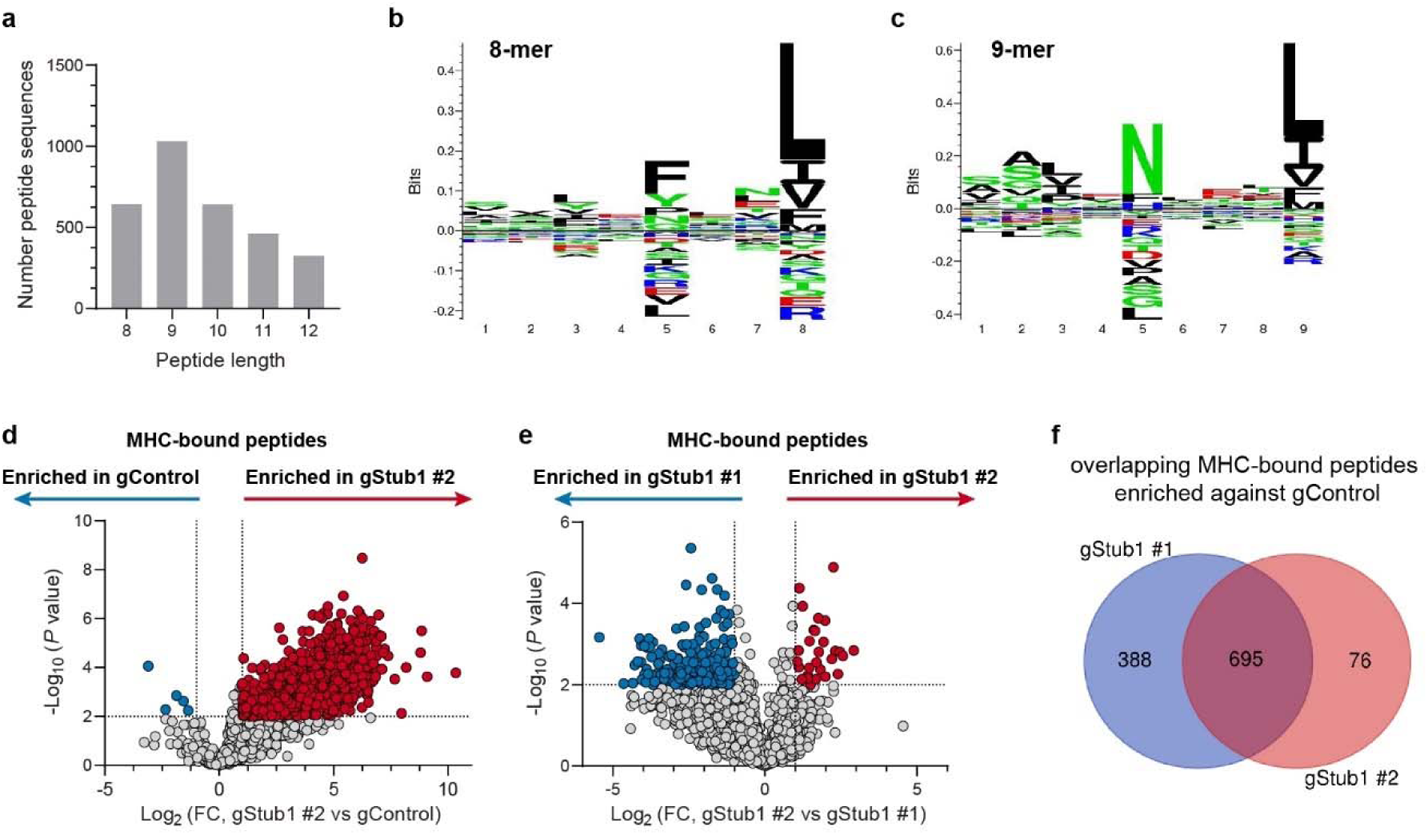
Immunopeptidomics of B16-F10 CRISPR cell lines. Related to Fig. 1g. a, Distribution of length for peptides identified by immunopeptidomics. b, c, Sequence motif of the 8- and 9-mers peptides identified by immunopeptidomics. d, e, Volcano plot showing differential presentation of MHC-associated peptide in the tumour cells, following stimulation with 0.10 ng ml^−1^ IFNγ for 24 hours. Red and blue circles highlight peptides significantly enriched in the respective tumour cells (2-fold cutoff, *P* ≤0.01; *n* = 3 biological replicates). f, Venn diagram showing the number of unique peptides enriched for gStub1 #1 (1,083 peptides) and gStub1 #2 (771 peptides) relative to the gControl cells. There are 695 MHC-bound peptides overlapping between the two clonal *Stub1*-null cells.

**Supplementary Fig. 3.**
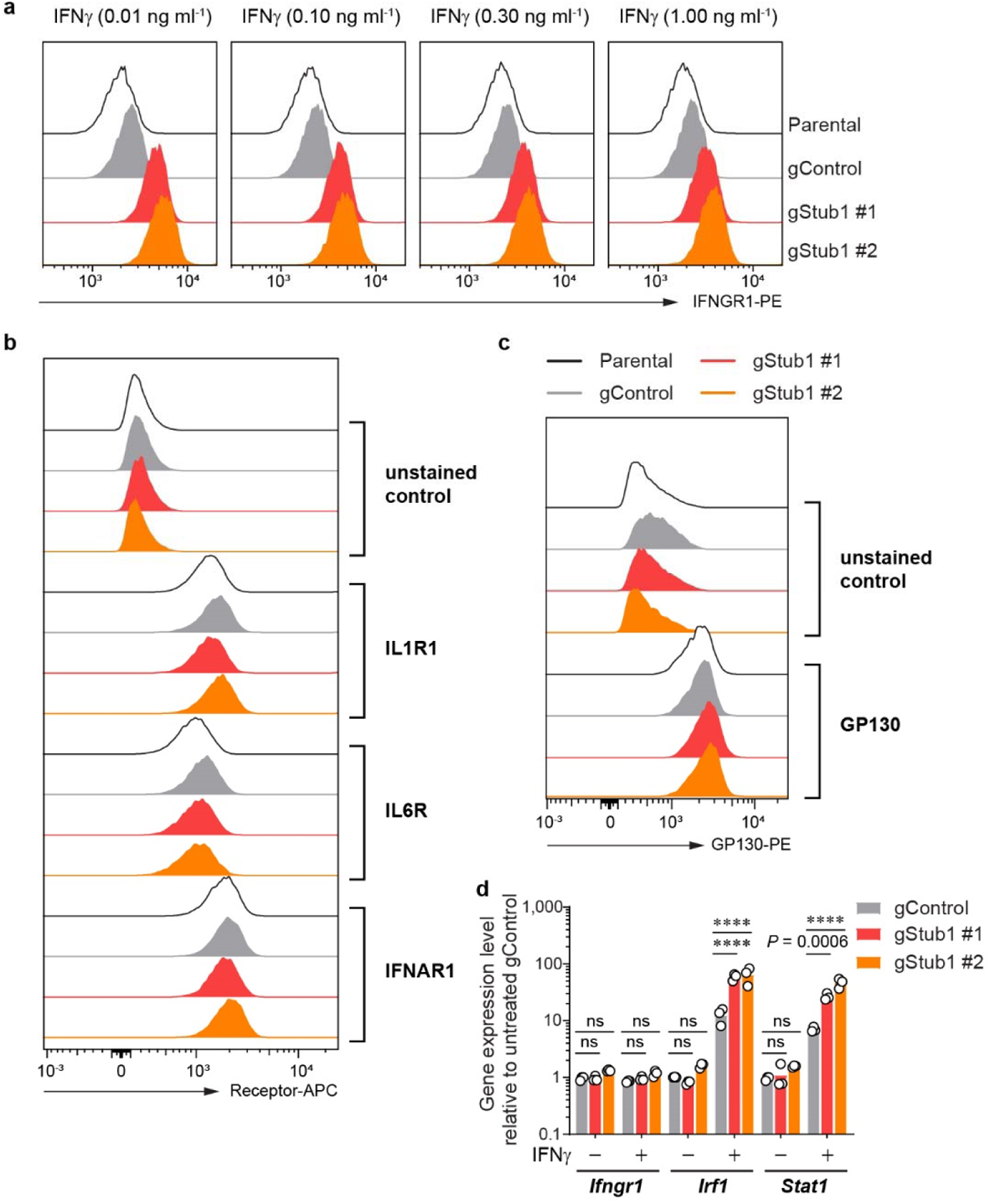
Analysis of the surface level of IFNGR1 and other immune-related receptors, and gene expression of *Ifngr1*, *Irf1* and *Stat1*. Related to Fig. 2. a, Flow cytometry analysis of cell surface IFNGR1 on parental, control or independent *Stub1*-null B16-F10 cells treated with IFNγ for 24 h. b, c, Flow cytometry analysis of cell surface IL1R1, IL6R or IFNAR1 (b) or GP130 (c) on parental, control or independent *Stub1*-null B16-F10 cells at resting state. d, qPCR analysis of the gene expression of *Ifngr1*, *Irf1* and *Stat1* relative to untreated gControl cells. Cells were stimulated with IFNγ (0.03 ng ml^−1^) for 6 h. Expression level was normalized to a reference gene (*Tbp*). Data are mean with all data points from three independent experiments (d). *P* values were determined by ordinary two-way ANOVA in each transcribed gene with Dunnett’s multiple comparisons test, **** *P* ≤0.0001, ns *P* >0.98 (d). Representative of three (a) or four (b–c) independent experiments.

**Supplementary Fig. 4.**
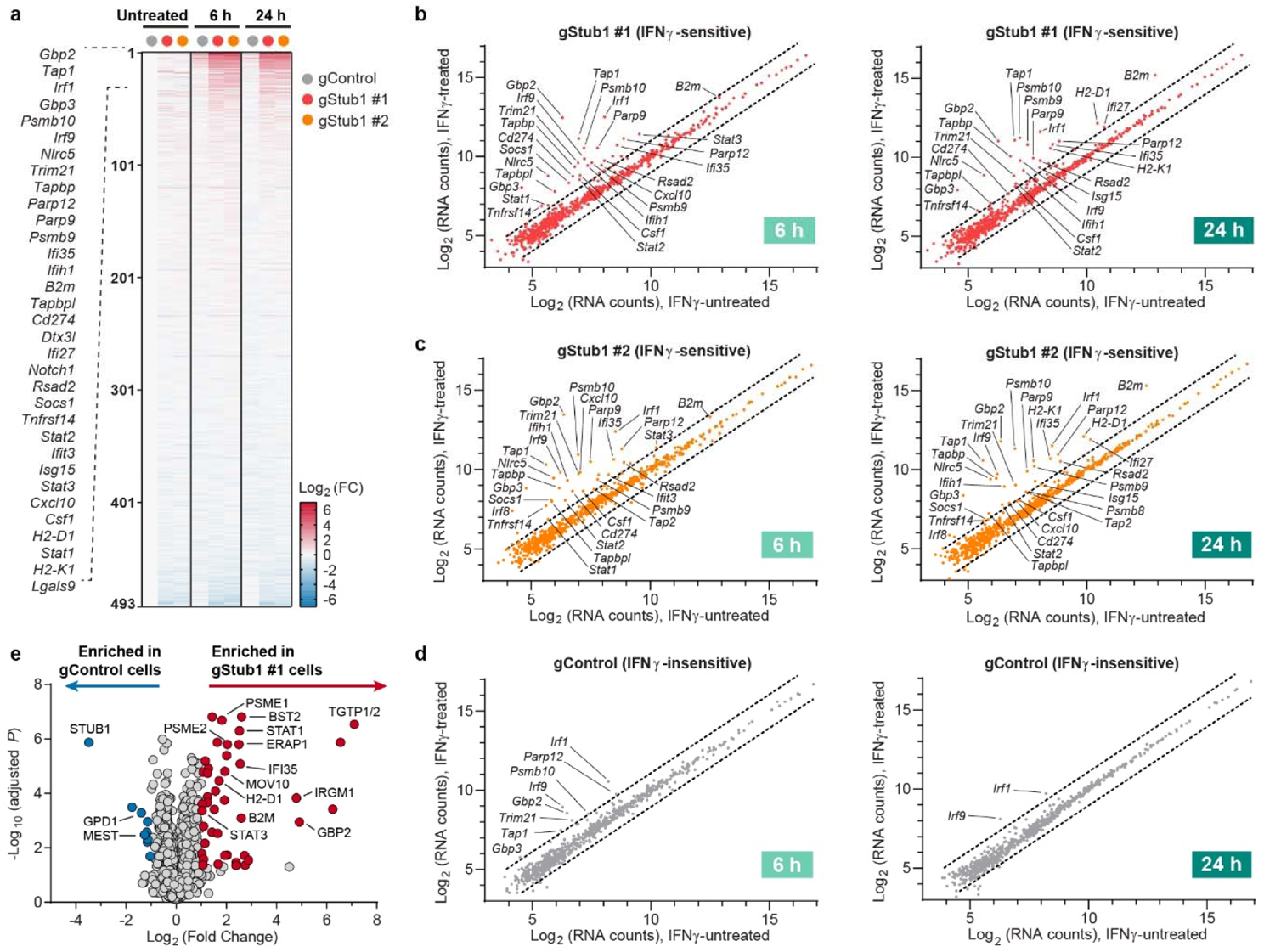
Loss of *Stub1* sensitized B16-F10 melanoma to IFNγ response as determined by gene expression profiling and proteome-wide studies. Related to Fig. 2. a, Heatmap showing relative gene expression profile of 493 out of 750 genes (NanoString PanCancer IO 360, Supplementary Table 3) in tumours cells which were either untreated or treated with 0.03 ng ml^−1^ IFNγ for 6 or 24 h. Weakly expressed genes (257 out of 750) were removed from the analysis. 33 genes upregulated by >2-fold in both *Stub1*-null cells relative to untreated gControl cells were listed explicitly (see Fig. 2e for the expanded heatmap). Refer to Method section for the description of fold change analysis. FC, fold change. b–d, Scatter plots showing the expression level of all 750 genes (Supplementary Table 3) in gControl cells (b) gStub1 #1 cells (c) or gStub1 #2 cells (d) before and after treatment with 0.03 ng ml^−1^ IFNγ for 6 h (left plot) or 24 h (right plot). Dotted lines depict the boundary of 2-fold change in the RNA counts which were normalized with 19 housekeeping genes. e, Volcano plot showing differential protein expression (Supplementary Table 4) in gStub1 #1 versus gControl cells, following treatment with 0.03 ng ml^−1^ IFNγ for 24 h. Red or blue circles highlight proteins being significantly enriched in gStub1 #1 or gControl cells respectively (2-fold cutoff, adjusted *P* ≤0.05; *n* = 6 replicates per cell group, 3 biological replicates × 2 MS replicates). Enriched proteins that overlap with gStub1 #2 cells (see Fig. 2f–g) are explicitly labeled in the plot. STUB1, GPD1 and MEST were consistently enriched in gControl cells as compared to either gStub1 #1 or gStub1 #2 cells (see Fig 2f). Statistics details were described in proteomics method section.

**Supplementary Fig. 5.**
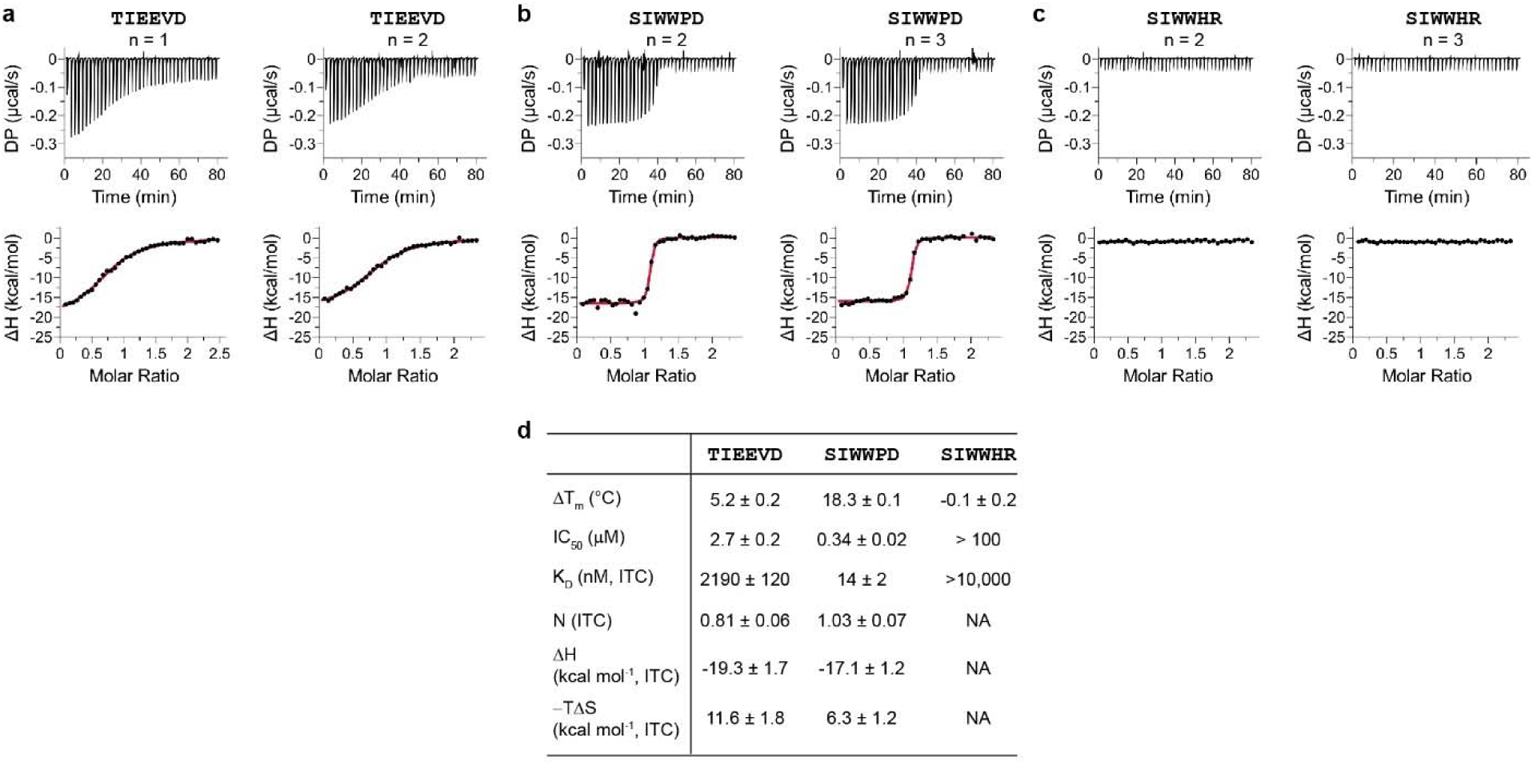
Validation of the binding of synthetic peptides to STUB1 with multiple biophysical assays. Related to Fig. 3. a–c, Binding of the synthetic peptides to STUB1 (aa25–aa153) as determined by isothermal titration calorimetry (ITC). The peptides contain free carboxylic acid at the C-terminus and are acetylated at the N-terminus. Positive control peptide (TIEEVD) is derived from the C-terminal end of HSPA8 – the endogenous binding substrate of STUB1 (a). SIWWPD is bound strongly to the protein (b), whereas SIWWHR is a non-binding control (c). d, Summarized results from all biophysical assays. The shift in the melting temperature (⊗T_m_) relative to the DMSO vehicle is reported as mean ± s.d. from three independent experiments. Half maximal inhibitory concentration (IC_50_) is reported as mean ± s.e. derived from the 4-parameter sigmoidal curve fitted with the data of six replicates derived from two independent fluorescence polarization experiments. Dissociation constant (K_D_), binding stoichiometry (N), enthalpy (ΔH) and entropy (−TΔS) are reported as mean ± s.d. from two (a) or three (b–c) independent ITC experiments.

**Supplementary Fig. 6.**
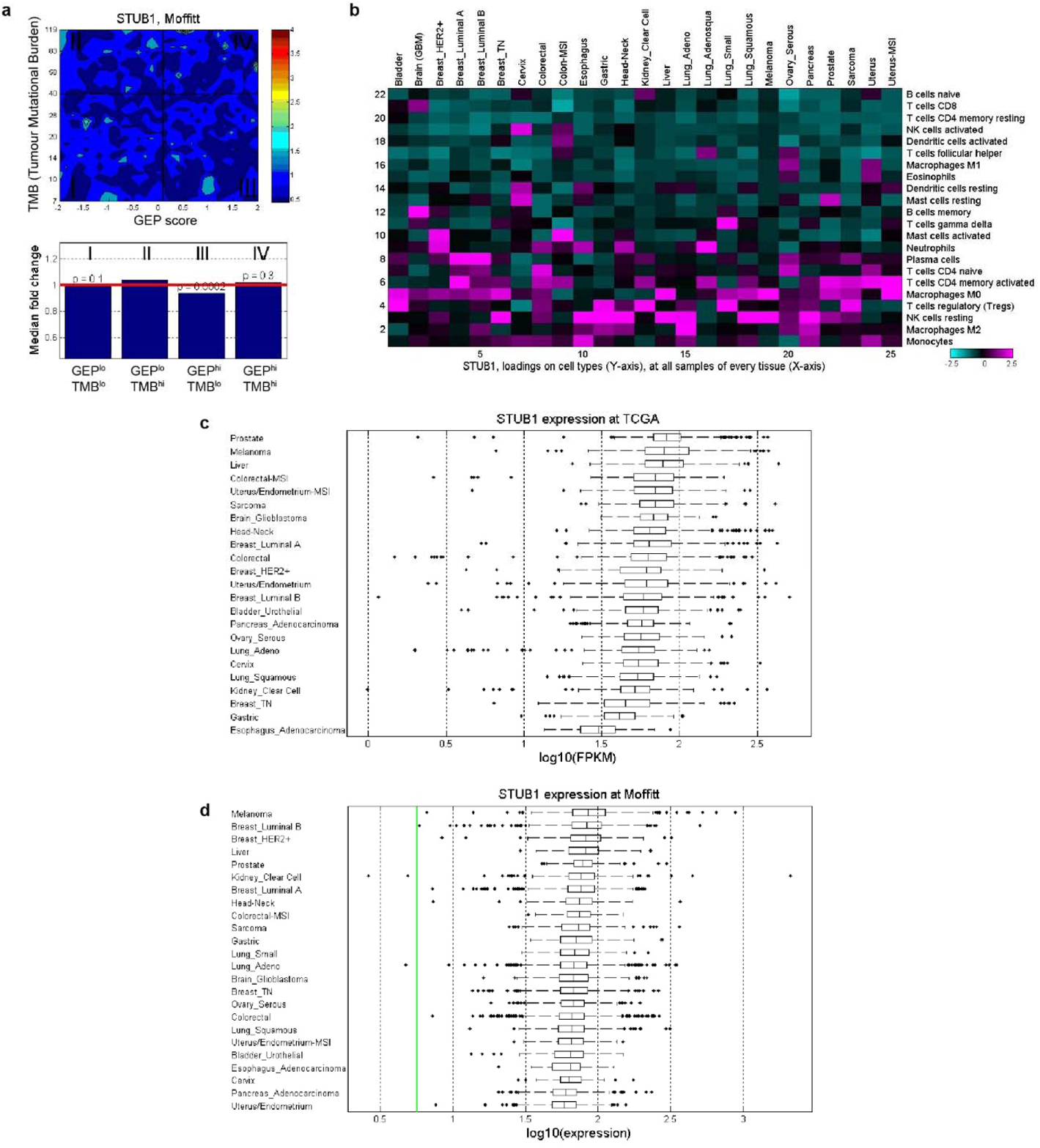
Correlation and expression of *STUB1* gene in Moffitt dataset. Related to Fig. 4. a, Contour plot illustrates the association of *STUB1* with TMB and GEP. Blue and red represent under- and overexpression, respectively. TMB cut-off was set at 40 and GEP cut-off corresponds to 55th percentile value for pan-cancer cohort. b, *In-silico* deconvolution analysis of bulk RNAseq data from Moffitt was used to establish the association between *STUB1* expression and different cell types. Deconvolution analysis, based on CIBERSORT, was performed separately for each tumor type. c, d, Relative *STUB1* expression level across major tumour tissues in TCGA (c) and Moffitt (d). Limit of detection >log_10_(−1.7) in TCGA. The green line depicts the limit of detection (d).

**Supplementary Fig. 7.**
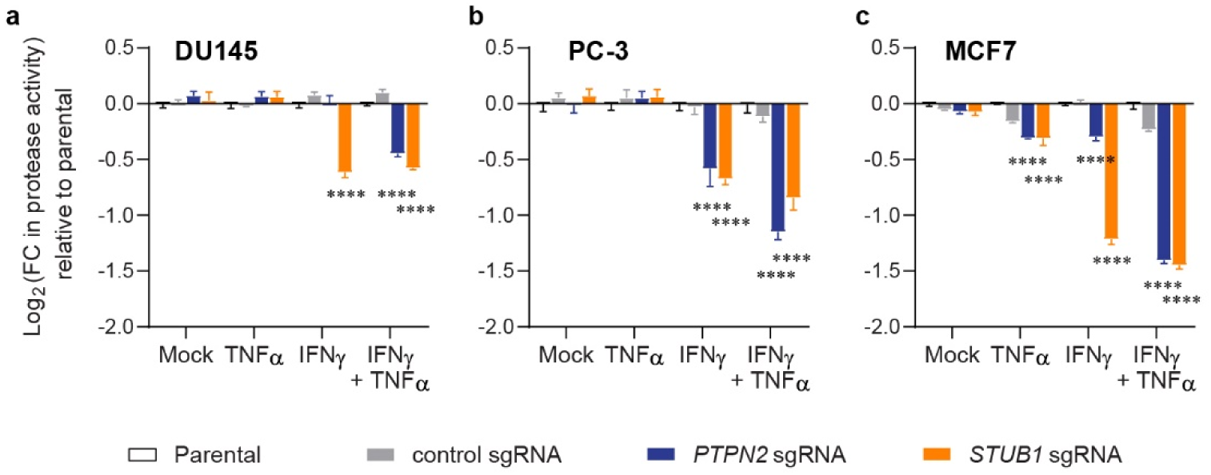
Inactivation of *STUB1* or *PTPN2* sensitized human tumour cells to growth inhibition induced by cytokines. Related to Fig. 5d. a–c, Fold change (FC) in live-cell protease activity relative to the parental cells as a quantification of viable cells. Measurements were performed using CellTiter-Fluor assay after 6-day treatment of DU145 (a), PC-3 (b), or MCF7 (c) cells and their corresponding CRISPR-edited lines with the cytokines (10 ng ml^−1^ each). Data are mean ± s.e.m. from three biological replicates (a–c). *P* values were determined by ordinary two-way ANOVA on Log2-transformed data with Dunnett’s multiple comparisons test versus parental cells, ** *P* ≤0.01, **** *P* ≤0.0001, ns *P* >0.90 (a–c).

**Supplementary Fig. 8.**
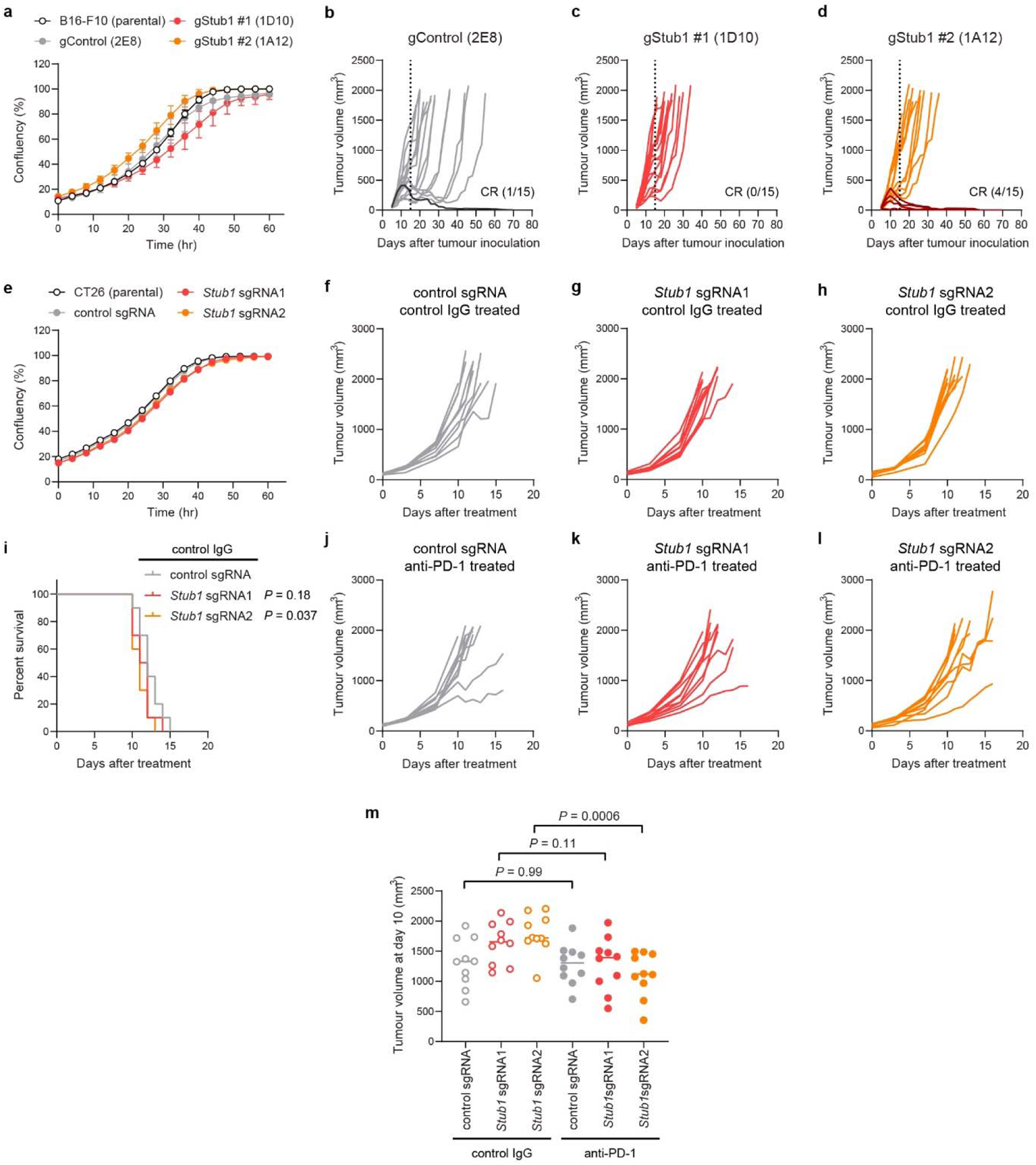
Additional data for the studies of syngeneic mouse models. Related to Fig. 6. a, Plot showing *in vitro* cellular growth kinetic of B16-F10 and the CRISPR clonal lines in standard 2D culture as measured by Incucyte (*n* = 3 biological replicates per cell type). b–d, Plot showing individual tumour volume of the CRISPR-edited B16-F10 clonal cells implanted into syngeneic mice (*n* = 15). CR, complete response. Dotted line indicates the day when mice received the last dose of anti-PD-1 antibody. e, Plot showing *in vitro* cellular growth kinetic of CT26 and the CRISPR lines in standard 2D culture as measured by Incucyte (*n* = 3 biological replicates per cell type). f–h, Plot showing individual tumour volume of the CRISPR-edited CT26 cells implanted into syngeneic mice (*n* = 10) treated with control antibody. i, Kaplan-Meier survival curves of tumour-bearing mice treated with control antibody. Median survival: control sgRNA, 12 days; *Stub1* sgRNA1, 11.5 days, *Stub1* sgRNA2, 11 days. j–l, Plot showing individual tumour volume of the CRISPR-edited CT26 cells implanted into syngeneic mice (*n* = 10) treated with anti-PD-1 antibody. m, Plot showing tumour volume at day 10 for the CRISPR-edited CT26 cells implanted into mice (*n* = 10) treated with either control or anti-PD-1 antibody. Representative of two independent experiments (a, e). *P* values were determined by Log-rank (Mantel-Cox) test versus control tumours (i). *P* values were determined by two-way ANOVA with Sidak’s multiple comparisons test (m). Data are mean with all data points derived from 10 mice per group (m).

**Supplementary Fig. 9.**
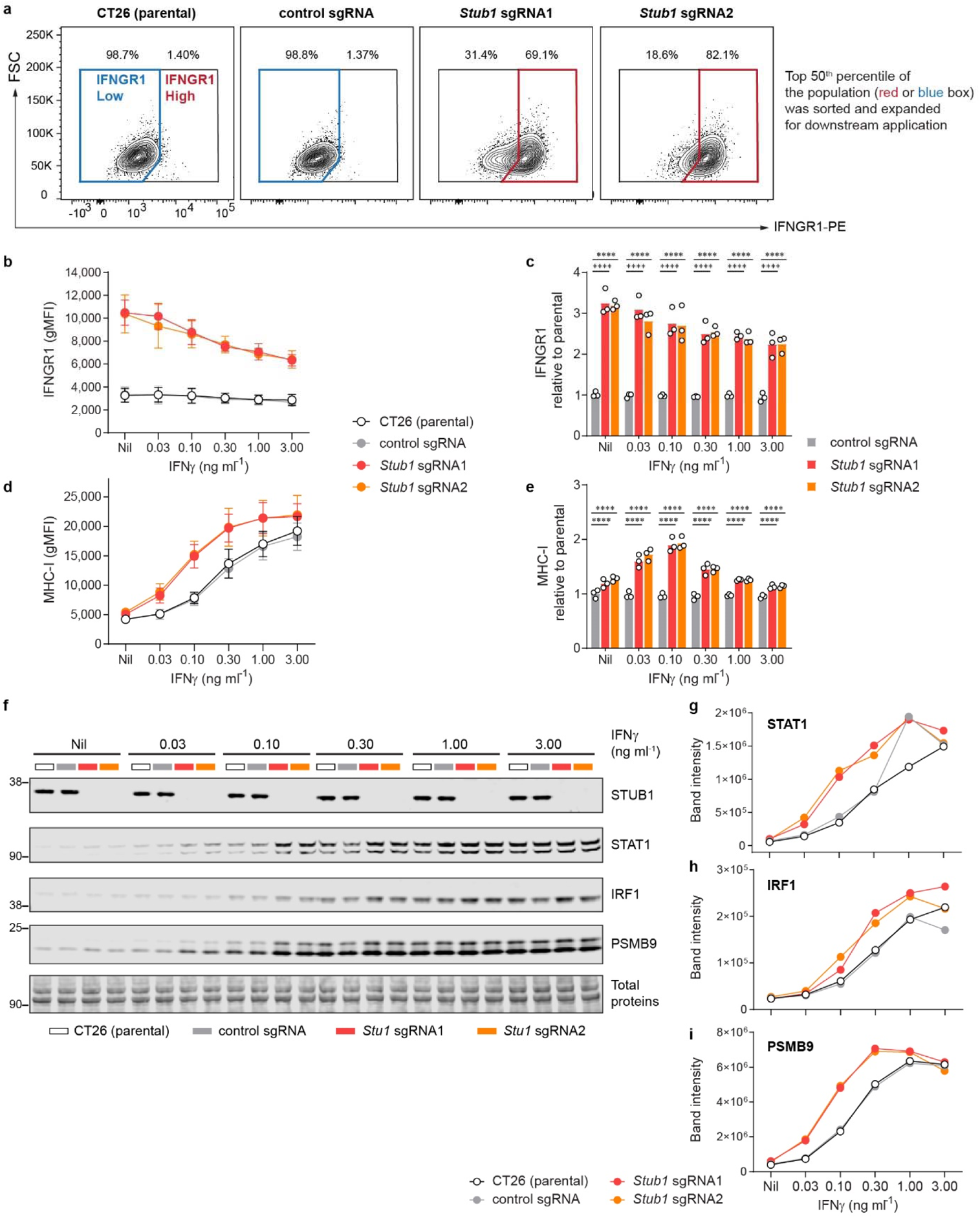
Loss of *Stub1* in CT26 tumour cells elevated the surface level of IFNGR1, leading to increased expression of MHC-I, STAT1, IRF1 and PSMB9 in response to IFNγ. Related to Fig. 6d–f and Supplementary Fig. 8e–m. a, Flow cytometry 2D plot showing the expression level of IFNGR1 on the cellular surface of CT26 and the corresponding CRISPR-edited cells. We gated for the population with high expression of IFNGR1 and sorted for the top 50^th^ percentile in the gated population (red box) to enrich the *Stub1*-null cells. Similar gating (IFNGR1 Low) and sorting (top 50^th^ percentile, blue box) strategies were consistently applied to the parental and control sgRNA targeting CT26 cells. b–e, Flow cytometry analysis of cell surface IFNGR1 (b, c) and MHC-I (d, e) expressed on parental, control or independent *Stub1*-null CT26 cells (sorted). gMFI, geometric mean fluorescence intensity. f–i, Western blot analysis of the expression level of STUB1, STAT1, IRF1, and PSMB9 in parental, control or independent *Stub1*-null CT26 cells (sorted). Band intensities were quantified with LI-COR Image Studio and normalized with total protein signal. The tumour cells were either untreated (Nil) or treated with recombinant mouse IFNγ for 24 h (b–i). Data are mean ± s.d. (b, d) or mean with all data points (c, e) from three independent experiments. *P* values were determined by ordinary two-way ANOVA on Log2-transformed data with Dunnett’s multiple comparisons test, **** *P* ≤0.0001 (c, e). Representative of two independent experiments (f–i).

